# The SWI/SNF subunit SWI3B functions with the m^6^A writer complex to establish embryo patterning in Arabidopsis

**DOI:** 10.64898/2026.08.13.744595

**Authors:** Wen Gong, Uwe Schwartz, Lifei Fu, Gernot Längst, Thomas Dresselhaus

## Abstract

N^6^-methyladenosine (m^6^A) is the most abundant mRNA modification in eukaryotes and is essential for Arabidopsis embryogenesis. However, how m^6^A mRNA methylation is coordinated with other regulatory pathways during development including embryogenesis remains largely unknown. Here, we report the SWI/SNF chromatin-remodeling subunit SWI3B as a *bona fide* interactor of the m^6^A methyltransferase MTA. Like m^6^A writer mutants, SWI3B is required for early embryo development. We demonstrate that the interaction between MTA and SWI3B is required for MTA function during embryogenesis. MTA and SWI3B are both required to establish the correct expression pattern of *WOX8* and proper auxin maxima during early embryogenesis. Transcriptome analysis of isolated embryos from *mta*, *swi3b*, and *fip37* mutants identified a shared set of upregulated transcripts, including STM as well as several NAC and ERF transcription factors that are normally absent or expressed at very low levels during early embryogenesis. Embryo-specific overexpression of *ANAC087* and *ERF114* genes phenocopied early embryonic defects observed in *mta* and *swi3b* mutants, indicating that their ectopic expression contributes to the observed developmental phenotype. Moreover, SWI3B and MTA are both required for m^6^A deposition on specific developmental transcripts. Together, our findings uncover a mechanism by which chromatin remodeling and m^6^A-mediated RNA regulation cooperate to suppress the precocious stability of key developmental regulators, thereby contributing to the establishment of the transcriptional program required for early embryo patterning in Arabidopsis.

**Highlights:**

- The SWI/SNF subunit SWI3B is a functional interactor of the m^6^A methyltransferase MTA during Arabidopsis embryogenesis
- SWI3B and MTA cooperate to establish embryo patterning, *WOX8* expression, and auxin maxima
- MTA and SWI3B suppress precocious expression of *STM*, *ANAC087* and *ERF114* transcription factors that disrupt early embryo development
- SWI3B links chromatin-associated regulation with m^6^A-mediated control of transcript stability

**Graphical abstract:** 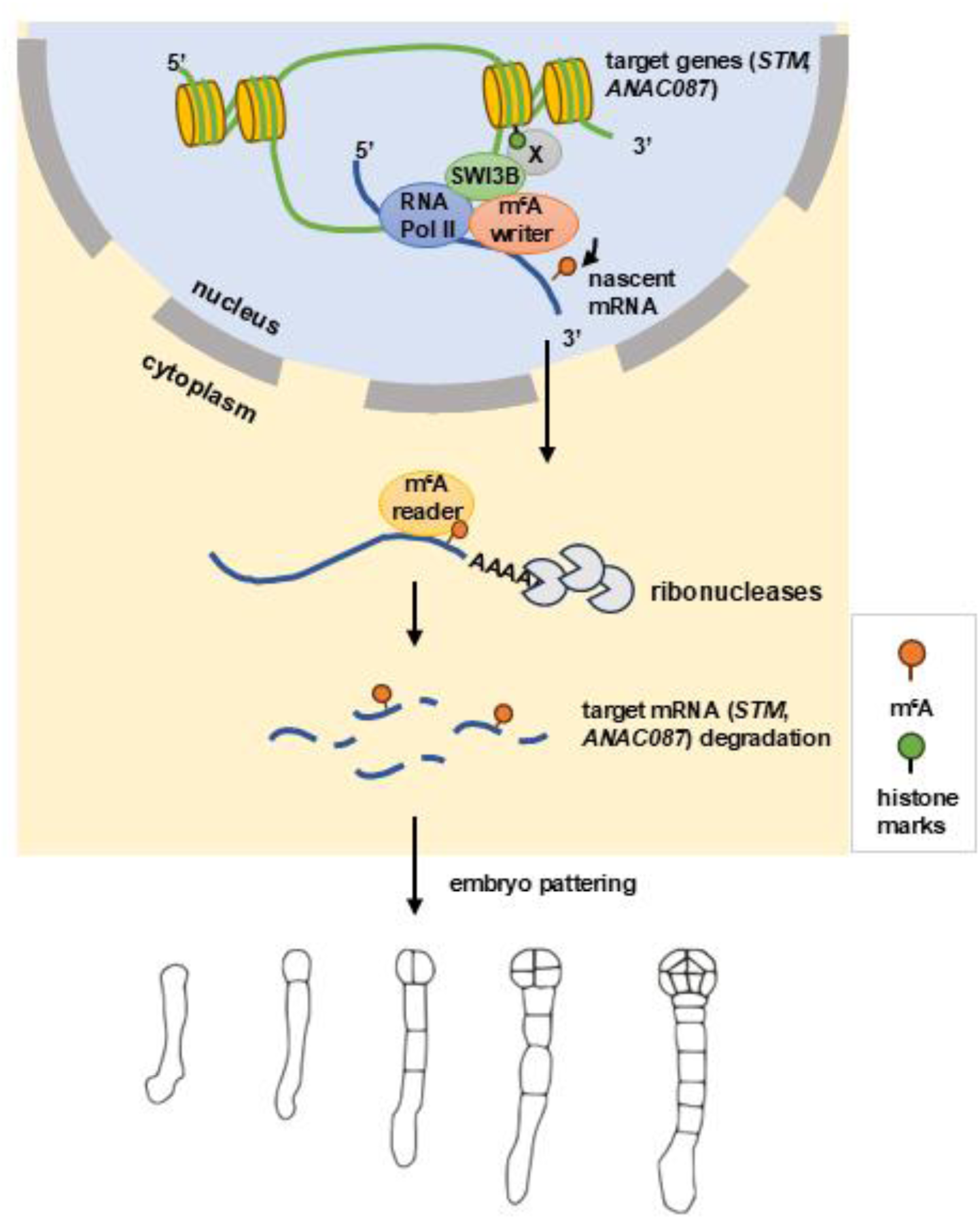

## Introduction

m^6^A is the most abundant internal modification of eukaryotic mRNA regulating gene expression at co- and post-transcriptional levels. The reversible m^6^A modification affects almost all aspects of RNA metabolism, including pre-mRNA splicing, mRNA export, mRNA stability and translational efficiency (Bhat, et al., 2020; Huang, et al., 2020; Reichel, et al., 2019; Yang, et al., 2018). m^6^A is deposited by the methyltransferase complex (m^6^A writers) pre-dominantly at the 3’ end of transcripts (Zaccara, et al., 2019; Anderson, et al., 2018; Meyer, et al., 2012). The m^6^A writer complex is formed by an assembly of proteins which have been characterized and described to be conserved between animals and plants. The mammalian m^6^A methyltransferase complex consists of methyltransferase-like 3 (METTL3) (Bokar, et al., 1997), methyltransferase-like 14 (METTL14) (Liu, et al., 2014), Wilms’ tumor 1-associating protein (WTAP) (Ping, et al., 2014), VIRMA (KIAA1429) (Schwartz, et al., 2014), RNA-binding motif protein 15 (RBM15) (Patil, et al., 2016), and zinc finger CCCH-type containing protein 13 (ZC3H13) (Wen, et al., 2018). While METTL3 and METTL14 constitute the m^6^A-METTL Complex (MAC), auxiliary proteins constitute the m^6^A-METTL Associated Complex (MACOM) (Knuckles, et al., 2018). In Arabidopsis, Methyltransferase A (MTA), the orthologue of METTL3 was identified as the catalytic component of m^6^A methyltransferase complex (Zhong, et al., 2008). FKBP12 interacting protein 37 kDa (FIP37, a homolog of WTAP) was the first identified member of the MACOM and was shown to be required for m^6^A modification in mRNA (Shen, et al., 2016; Zhong, et al., 2008). The other component of MAC, Methyltransferase B (MTB, the orthologue of METTL14), together with the other components of the MACOM, Virilizer (VIR, the orthologue of VIRMA) and HAKAI (the orthologue of an E3 ubiquitin ligase) were identified (Ruzicka, et al., 2017). The Arabidopsis homolog of RBM15 is FLOWERING LOCUS PA (FPA), which was shown to co-purified with m^6^A writer proteins (Parker, et al., 2021). Later, the HAKAI-interacting zinc finger protein 1 and 2 (HIZ1 and HIZ2) were identified as members of the MACOM, which HIZ2 is likely the plant homolog of ZC3H13 (Zhang, et al., 2022). Null mutants of MTA or any other members of the core writer complex (MTB, FIP37 and VIR) are embryo-lethal, indicating an essential function of m^6^A mRNA modification during embryogenesis (Ruzicka, et al., 2017; Shen, et al., 2016; Zhong, et al., 2008). Defects in embryo development of null mutants of MTA and FIP37 can be complemented by using the embryo-specific promoter *ABSCISIC ACID INSENSITIVE 3* (*ABI3*) and *LEAFY COTYLEDON 1* (*LEC1*) respectively (Shen, et al., 2016; Bodi, et al., 2012). FIP37 is essential for preventing shoot apical meristem over proliferation by mediating the m^6^A mRNA modification in the mRNA for two stem cell regulators, WUSCHEL (WUS) and SHOOT MERISTEMLESS (STM), resulting in a shorter lifetime of the mRNAs (Shen, et al., 2016). These findings established a molecular link between m^6^A mRNA modification and regulation of gene expression spatially and temporally during plant development. Despite the indispensable roles of m^6^A writers in plant embryo development, so far, the genes and pathways mediated by m^6^A mRNA modification during embryogenesis remain unknown.

Further research demonstrated that there are interconnections between m^6^A modification and histone modification, which has been demonstrated so far exclusively in animals (Shen and Yu, 2025; Hofler and Duss, 2024; Hu, et al., 2024; Xu, et al., 2022). In mammalian cells, histone H3 trimethylation at lysine 36 (H3K36me3) is bound directly by METTL14, which in turn facilitates m^6^A modification of actively transcribed RNAs (Huang, et al., 2019). METTL3 gets recruited to repressive histone marks H3K9m3 and H4K20m3, and the interaction is maintained by m^6^A reader YTHDC1 (Xu, et al., 2021). On the other hand, several studies demonstrate that m^6^A influences histone modification. Direct interaction between METTL14 and H3K27me3 leads to the recruitment of histone demethylase KDM6B, which removes H3K27me3 (Dou, et al., 2023). The interplay between m^6^A modification and histone regulation remains poorly understood in plants. Emerging evidence suggests a potential connection between these two epigenetic layers. For example, ectopic expression of the human m^6^A demethylase FTO in rice and potato alters histone methylation patterns while enhancing plant growth and yield (Yu, et al., 2021). In addition, H3K36me2 has been linked to m^6^A deposition on mRNAs, suggesting that histone modifications may influence m^6^A establishment. Conversely, m^6^A-marked transcripts could recruit factors that modulate chromatin states, although the underlying mechanisms remain unknown. Together, these observations point to a functional crosstalk between RNA methylation and histone modification in plants, but its molecular basis and developmental significance remain largely unexplored.

In this study, we identify the SWI/SNF chromatin-remodeling subunit SWI3B as a functional partner of the m^6^A methyltransferase MTA and demonstrate that their interaction is essential for establishing the transcriptional program underlying early embryo patterning. Embryonic transcriptomic analysis reveals a potential shared set of target transcripts of MTA, FIP37 and SWI3B, including several developmental associated transcription factors. We further demonstrate that MTA and SWI3B mediate m^6^A mRNA modification on two transcription factor genes and negatively correlate with their mRNA stability. Our findings reveal a previously unrecognized mechanism linking chromatin remodeling with m^6^A-dependent post-transcriptional regulation in Arabidopsis.

## Results

### MTA and FIP37 are required for early embryo patterning

To study the function of m^6^A writers MTA and FIP37 during Arabidopsis embryo development, we characterized a T-DNA insertion mutant in *MTA*, which we termed *mta-2* (SALK_123823) and a previously reported T-DNA insertion mutant in *FIP37*, *fip37-4* (SALK_018636) (Figure S1A). Homozygous plants containing these two insertions could not be identified from self-pollinated heterozygous mutant plants. About one-quarter of the seeds of *mta-2/+* and *fip37- 4/+* were white in color, while the rest containing green colored bent-cotyledon stage embryo (Figure 1A, 1B), indicating a homozygous embryo-lethal phenotype, which is consistent with previous findings (Shen, et al., 2016; Zhong, et al., 2008). To investigate at which developmental stage the defects occur, the segregating embryos from *mta-2/+* and *fip37-4/+* were analyzed. In WT embryos at the 8-cell stage, the basal cell divided transversely, forming the suspensor. However, about one-quarter of embryos from *mta-2/+* and *fip37-4/+* showed abnormal longitudinal cell division pattern in the suspensor (Figure 1D). At the globular stage, the hypophysis in WT embryos divided asymmetrically to produce an upper lens-shaped cell and a lower cell connecting the embryo proper to the suspensor. In contrast, defective embryos from *mta-2/+* and *fip37-4/+* exhibited abnormal longitudinal division of the hypophysis, accompanied by irregular morphology of the embryo proper. In *mta-2/+*, a severe thread-like embryo phenotype was observed, characterized by the absence of periclinal cell division in the embryo proper and abnormal cell divisions in the suspensor (Figure 1E). To determine the developmental stage at which the mutant embryos became arrested, white ovules containing arrested embryos were examined. Arrested embryos from *mta-2/+* contained an irregular globular-like embryo proper, and some also displayed abnormal cell divisions in the suspensor, resulting in an embryo proper-like structure (Figure S1C). In contrast, defective embryos from *fip37-4/+* developed to the transition stage, and subsequently arrested between the heart and torpedo stages, exhibiting an abnormal single-cotyledon phenotype (Figure S1B, S1C). To examine the expression pattern of *MTA* and *FIP37*, genomic constructs carrying a C-terminal GFP fusion were generated. Homozygous *pMTA:MTA-GFP mta-2* and *pFIP37:FIP37-GFP fip37-4* transgenic lines were identified (Figure 1A), indicating that both GFP fusion proteins are functional. Accordingly, the frequencies of embryo patterning defects and embryo arrest were reduced to negligible levels in both transgenic lines (Figure 1B, 1C, S1D). Both, *pMTA:MTA-GFP* and *pFIP37:FIP37-GFP* were expressed in female and male germline cells, and in embryos throughout embryogenesis, exhibiting an ubiquitous expression pattern. MTA-GFP and FIP37-GFP were exclusively localized to nuclei (Figure 1F, 1G, S2).

**Figure 1.**
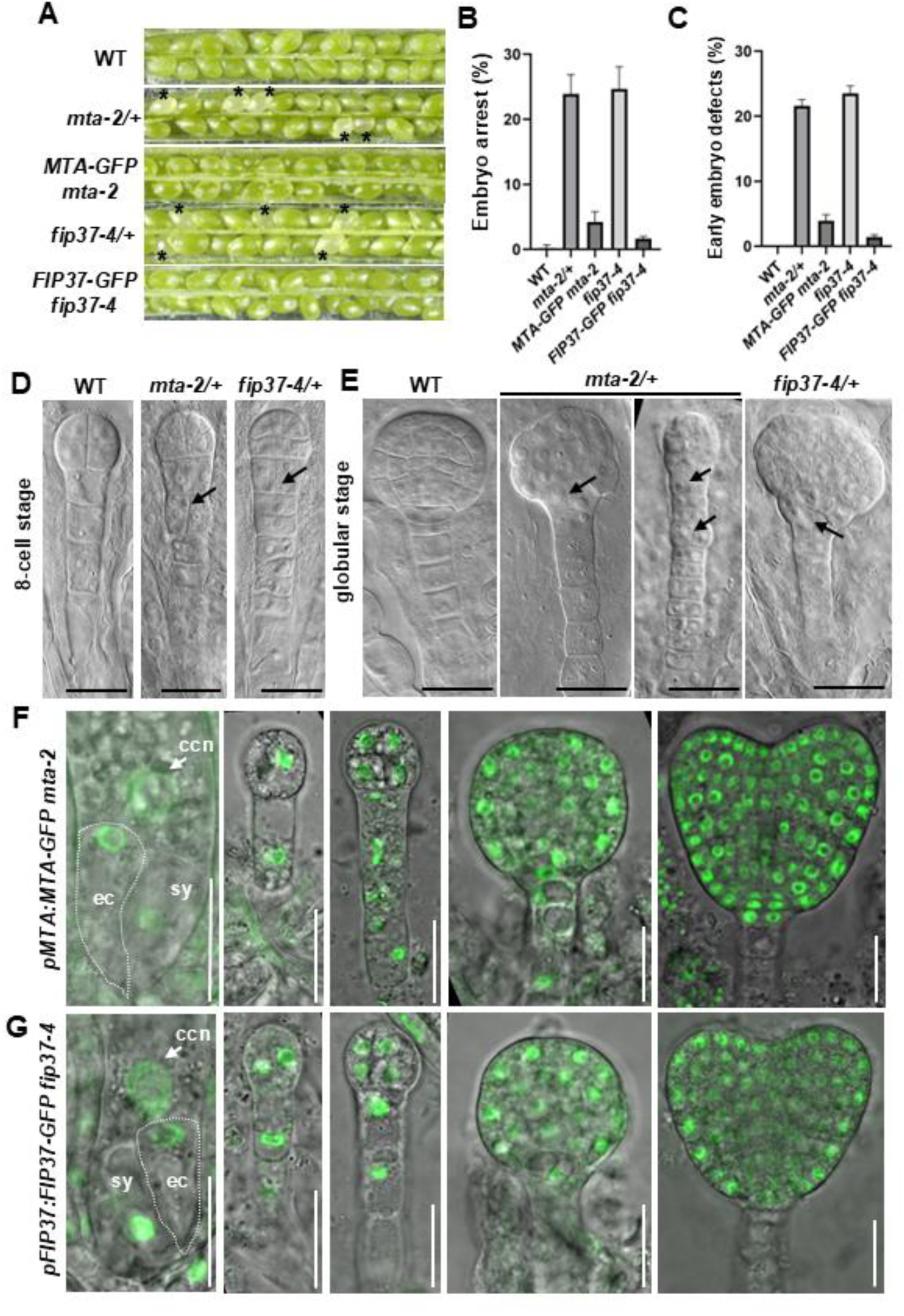
MTA and FIP37 are required for early embryo patterning. (A) Dissected siliques of the indicated genotypes. Asterisks indicate white, aborted ovules. (B–C) Quantification of the percentages of embryo arrest (B) and embryo patterning defects at the 8-cell stage (C) in the indicated genotypes. Data are presented as mean ± SD from three biological replicates. (D–E) DIC microscopy of 8-cell-stage (D) and globular-stage (E) embryos of WT, *mta-2/+*, and *fip37-4/+*. Arrows indicate abnormal cell division planes. Scale bars, 20 μm. (F) Expression pattern of *pMTA:MTA-GFP* in the *mta-2* background in unfertilized ovules and embryos from the 1/2-cell to heart stage. (G) Expression pattern of *pFIP37:FIP37-GFP* in the *fip37-4* background in unfertilized ovules and embryos from the 2-cell to heart stage. Abbreviations: ccn, central cell nucleus; ec, egg cell; sy, synergid cell. Scale bars, 20 μm.

### SWI3B physically interacts with MTA

Previous studies have established extensive crosstalk between m^6^A modification and histone modifications, yet the molecular mechanisms linking the m^6^A writer complex to the chromatin-remodeling machinery remain largely unexplored. It is proposed that the histone marks and an unidentified protein recruit m^6^A writers to the target mRNA, leading to deposition of the m^6^A mark (Hu, et al., 2024; Yu, et al., 2021; Shim, et al., 2020). To explore potential links between the m^6^A writer complex and chromatin regulation, we previously screened chromatin-remodeling factors and histone-modifying enzymes for interaction with MTA using bimolecular fluorescence complementation (BiFC). This screen identified the SWI/SNF chromatin-remodeling subunit SWI3B as a candidate MTA-interacting protein. Reconstituted YFP fluorescence was detected in the nuclei of Arabidopsis protoplasts co-expressing *MTA-nYFP* and *SWI3B-cYFP*, indicating that MTA interacts with SWI3B. As expected, MTA also interacted with its known partner FIP37, whereas an interaction was not detected between MTA and GIF2 (a subunit of SWI/SNF complex), which served as a negative control. In addition, we could show that SWI3B interacted also with FIP37. The BiFC signals were confined to the nucleus and were enriched in discrete granule-like foci (Figure 2A). To validate the interaction *in vivo*, co-immunoprecipitation (Co-IP) assays were performed using transgenic plants expressing *pMTA:MTA-GFP* and *pSWI3B:SWI3B-FLAG*. Immunoprecipitation of MTA-GFP efficiently co-precipitated SWI3B-FLAG, confirming the interaction between MTA and SWI3B (Figure 2B). An interaction was not detected between MTA and GIF2 (a component of SWI/SNF complex) (Figures 2B). In contrast to the BiFC assay, an interaction between SWI3B and FIP37 was not detected by Co-IP (Figure 2C). To further define the MTA regions involved in SWI3B interaction, we used AlphaFold3 (Abramson, et al., 2024) to predict the structure of a complex comprising MTA, MTB, FIP37, SWI3B, and a target RNA. AlphaFold3 generated a model with high overall confidence (Figure 2D). The model predicted multiple contacts between MTA and SWI3B, including electrostatic interactions between amino acids K443 and R446 of MTA and D64 and E278 of SWI3B, respectively (Figures 2E, 2F). Notably, amino acid D64 is located within the SWIRM domain of SWI3B, which is known to mediate protein–protein interactions during the assembly of chromatin-associated protein complexes (Aravind and Iyer, 2002). The model further predicted that amino acid residues Q334, E340, and R347 of MTA interact with Q208, R220, and E227 within the WTAP domain of FIP37 (Figure S3A). Several residues within the MT-A70 methyltransferase domain of MTA were predicted to contact the target RNA (Figure S3A).

**Figure 2.**
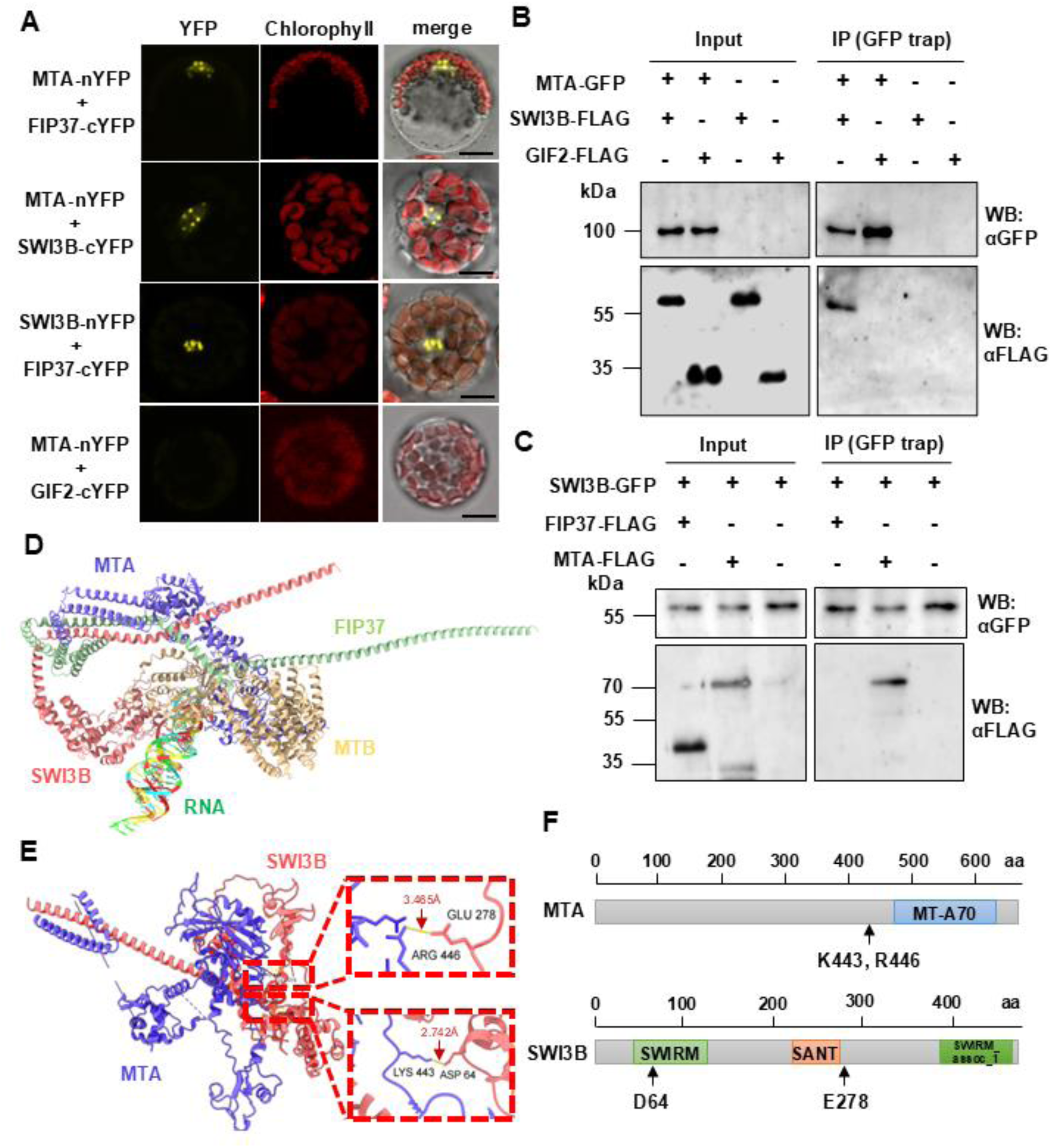
SWI3B physically interacts with MTA. (A) BiFC assays in Arabidopsis leaf mesophyll protoplasts showing interactions between MTA-nYFP and SWI3B-cYFP, and between SWI3B-nYFP and FIP37-cYFP. MTA-nYFP with FIP37-cYFP serves as a positive control, and MTA-nYFP with GIF2-cYFP serves as a negative control. Scale bars, 10 μm. (B) Co-IP assays using 5-day-old transgenic seedlings expressing *pMTA:MTA-GFP*/*pSWI3B:SWI3B-FLAG*, *pMTA:MTA-GFP*/*pGIF2:GIF2-FLAG*, *pSWI3B:SWI3B-FLAG* alone, or *pGIF2:GIF2-FLAG* alone. Immunoblots probed with anti-GFP and anti-FLAG antibodies are shown. (C) Co-IP assays using 5-day-old transgenic seedlings expressing *pSWI3B:SWI3B-GFP*/*pFIP37:FIP37-FLAG*, *pSWI3B:SWI3B-GFP*/*pMTA:MTA-FLAG*, or *pSWI3B:SWI3B-GFP* alone. Immunoblots probed with anti-GFP and anti-FLAG antibodies are shown. (D) AlphaFold3 model of a complex comprising MTA (blue), MTB (yellow), FIP37 (green), SWI3B (red), and a 95-nt RNA sequence surrounding the *ACTB* stop codon and two m^6^A modification sites (Tang, et al., 2025). The ribonucleotides of the RNA target are shown in different colors. (E) AlphaFold3 model of the predicted interaction between MTA (blue) and SWI3B (red). Enlarged regions outlined by red dashed boxes show the predicted interacting residues in MTA and SWI3B. (F) Schematic representation of MTA and SWI3B proteins. Amino acid positions are indicated above schematics, and colored boxes denote functional domains. Arrows indicate residues predicted to mediate the MTA-SWI3B interaction.

### SWI3B is required for early embryo development and colocalizes with m^6^A writers

Next, we investigated whether SWI3B is also required for embryo development like m^6^A writers. A previous study identified two T-DNA insertion mutant alleles in the gene *SWI3B*, named *swi3b-1* and *swi3b-2*. Both alleles were reported to be homozygous embryo lethal (Sarnowski, et al., 2005). To determine the stage of embryogenesis at which the defects arise, we obtained the T-DNA mutant *swi3b-2*/+ for further analysis (Fig S4A). To test whether SWI3B plays a role for early embryo patterning, we performed DIC microscopy of cleared ovules. In the *swi3b-2/+* mutant, 24.6±0.8% of segregating embryos exhibited abnormal cell division patterns (anticlinal cell division in the embryo proper and longitudinal division of the hypophysis) from 8-cell to 16-cell embryo stage, instead of the periclinal cell division to form the protoderm and asymmetric cell division of the hypophysis in WT (Figure 3A, 3C). At the globular stage, defective embryos from *swi3b-2/+* exhibited additional abnormal division patterns of the hypophysis, accompanied by irregular morphology of the embryo proper, which resemble *mta-2/+* and *fip37-4/+* (Figure 3B). Subsequently, about one-quarter of embryos became arrested (Figure 3D). Arrested embryos from *swi3b-2/+* contained an irregular globular-like embryo proper (Figure S4B). To examine the expression pattern of *SWI3B*, a genomic construct carrying a C-terminal GFP fusion was generated. The *pSWI3B:SWI3B-GFP swi3b-2* line fully complemented the embryo patterning defects and embryo lethality of *swi3b-2* (Figure 3C, 3D), indicating that the GFP fusion protein is functional. *pSWI3B:SWI3B-GFP* was expressed in the egg cell and, following fertilization, throughout embryogenesis. *SWI3B-GFP* was ubiquitously expressed until the globular stage, after which its expression became enriched in the epidermis and the shoot apical meristem region from the transition stage onward (Figure 3E). To determine whether SWI3B co-localizes with m^6^A writers during embryogenesis, *pSWI3B:SWI3B-mScarlet* was crossed with *pMTA:MTA-GFP* and *pFIP37:FIP37-GFP*, respectively. SWI3B-mScarlet extensively co-localized with both, MTA-GFP and FIP37-GFP in the nucleus from the 2-cell to globular embryo stage (Figure 3F, 3G and S3). Notably, all three proteins were enriched in discrete nuclear granule-like foci, consistent with previous reports that the m^6^A writer complex is enriched in nuclear speckles involved in RNA processing and alternative splicing (Ping, et al., 2014).

**Figure 3.**
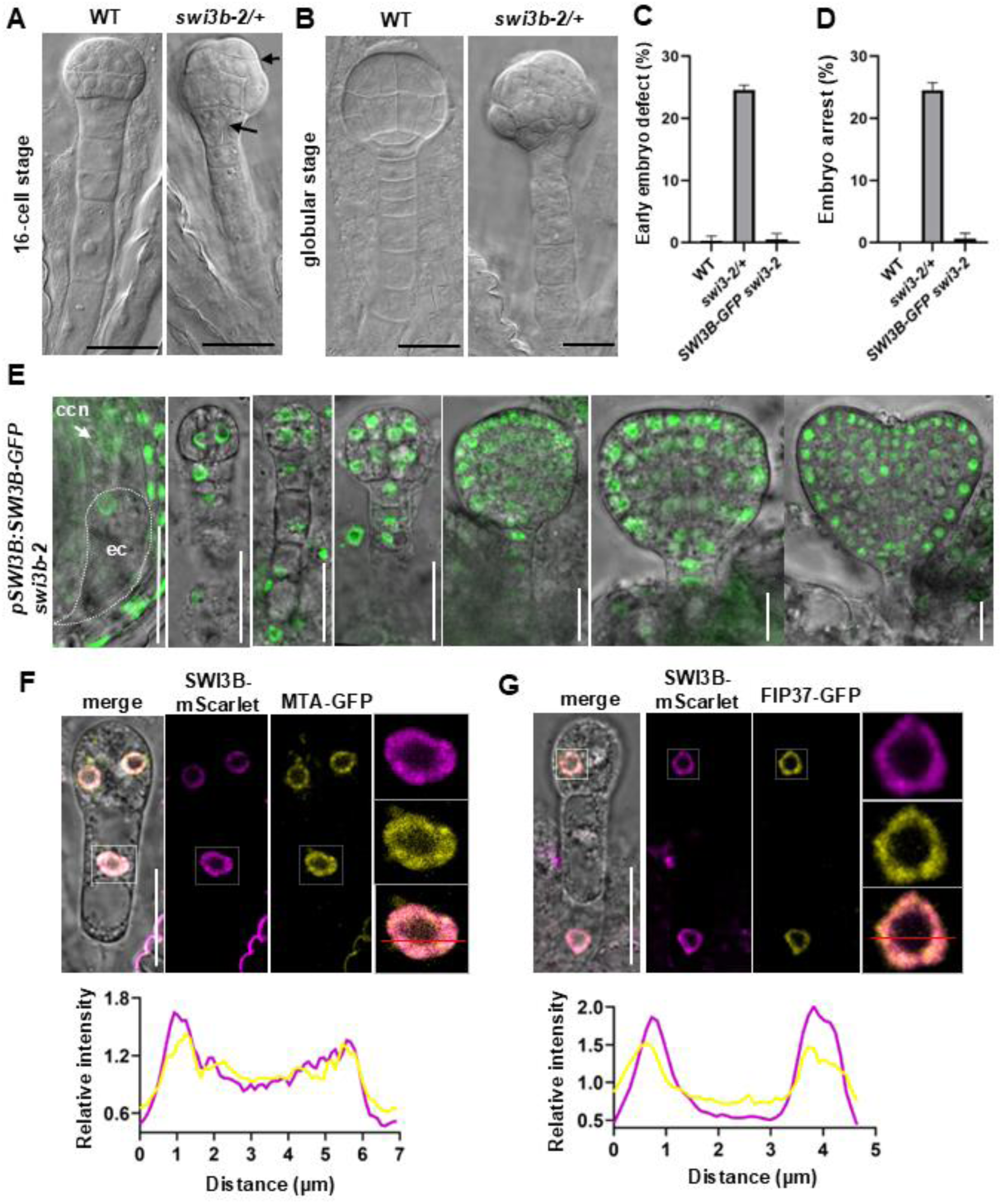
SWI3B is required for early embryo development and colocalizes with m^6^A writers. (A–B) DIC microscopy of 16-cell-stage (A) and globular-stage (B) embryos of WT and *swi3b-2/+*. Arrows indicate abnormal cell division planes. Scale bars, 20 μm. (C) Quantification of the percentage of embryos exhibiting patterning defects at the 8- to 16-cell stage. Data are presented as mean ± SD from three biological replicates. (D) Quantification of the percentages of embryo arrest in indicated genotypes. Data are presented as mean ± SD from three biological replicates. (E) Expression pattern of *pSWI3B:SWI3B-GFP* in the *swi3b-2* background in unfertilized ovules and embryos from the 2-cell to heart stage. Abbreviations: ccn, central cell nucleus; ec, egg cell. Scale bars, 20 μm. (F) Colocalization of MTA-GFP and SWI3B-mScarlet at the 2-cell-stage embryo. The region outlined by white dashed lines is enlarged in the right panels. Fluorescence intensity profiles of mScarlet and GFP along the indicated red line are shown. (G) Colocalization of FIP37-GFP and SWI3B-mScarlet at the 2-cell-stage embryo. The region outlined by white dashed lines is enlarged in the right panels. Fluorescence intensity profiles of mScarlet and GFP along indicated red line are shown.

### The interaction of MTA with SWI3B is required for MTA function during embryo development

To determine whether the interaction with SWI3B is required for MTA function during embryogenesis, we generated an MTA mutant in which amino acid residues K443 and R446 that were predicted by AlphaFold3 to mediate the interaction with SWI3B, were substituted with glutamic acid (mMTA; Figure 4A). In BiFC assays, reconstituted YFP fluorescence was readily detected in the nuclei of Arabidopsis protoplasts co-expressing MTA-nYFP and SWI3B-cYFP, whereas fluorescence was not observed between mMTA and SWI3B. This indicates that the mutations abolished the interaction. In contrast, mMTA retained its interaction with FIP37 (Figure 4B).

**Figure 4.**
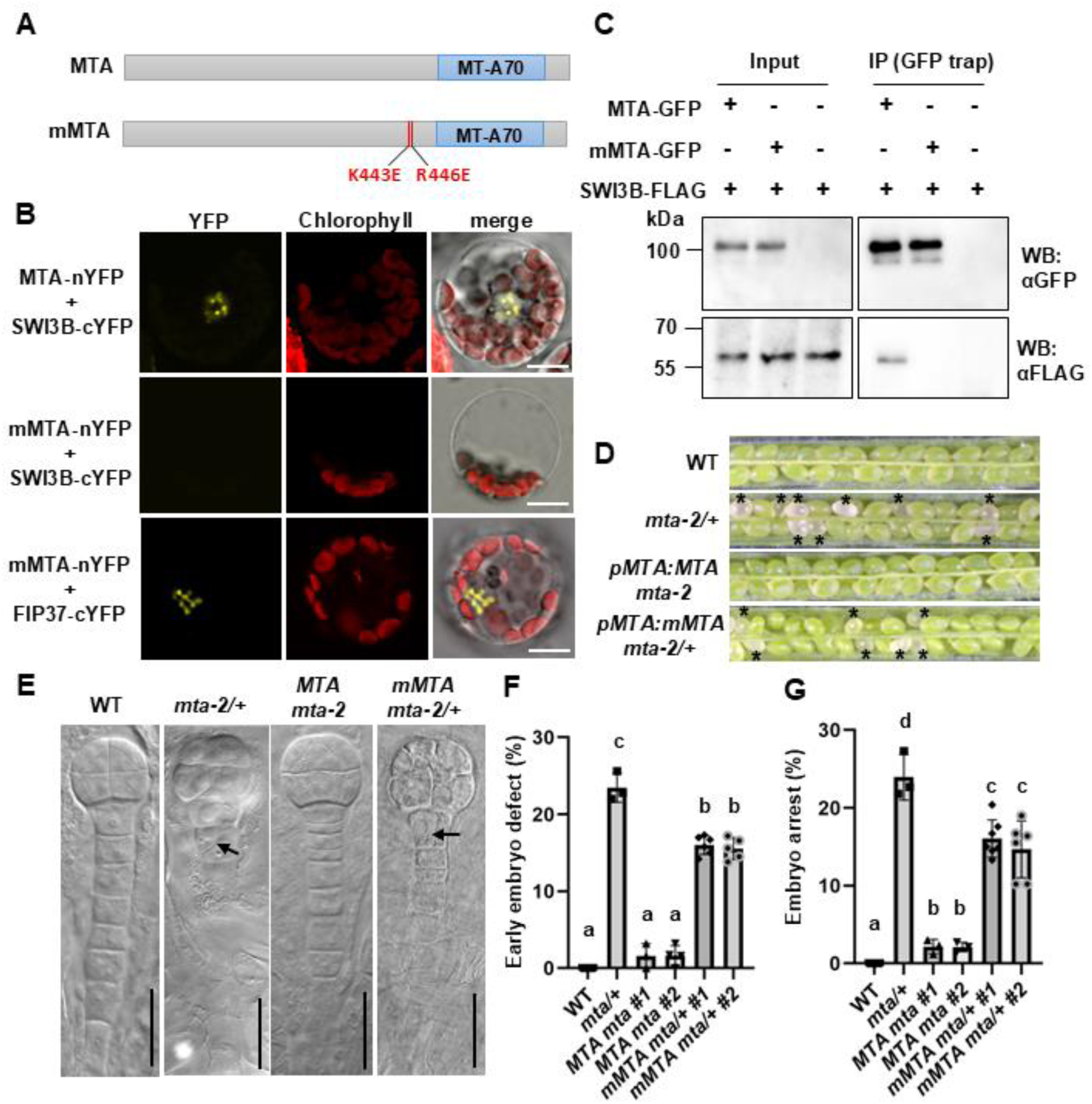
The interaction of MTA with SWI3B is required for its function in embryo development. (A) Schematic representation of MTA and mutated MTA (mMTA) proteins. K443E and R446E amino acid substitutions are indicated by red lines. (B) BiFC assays in *Arabidopsis* leaf mesophyll protoplasts showing loss of interaction between mMTA-nYFP and SWI3B-cYFP. MTA-nYFP with SWI3B-cYFP serves as a positive control. The interaction between mMTA-nYFP and FIP37-cYFP is retained. Scale bars, 20 μm. (C) Co-IP assays using 5-day-old transgenic seedlings expressing *pMTA:MTA-GFP*/*pSWI3B:SWI3B-FLAG*, *pMTA:mMTA-GFP*/*pSWI3B:SWI3B-FLAG*, or *pSWI3B:SWI3B-FLAG* alone. Immunoblots probed with anti-GFP and anti-FLAG antibodies are shown. (D) Dissected siliques of indicated genotypes. Asterisks indicate white, aborted ovules. (E) DIC microscopy of 16-cell-stage embryos of WT, *mta-2/+*, *pMTA:MTA mta-2*, and *pMTA:mMTA mta-2/+*. Arrows indicate abnormal cell division planes. Scale bars, 20 μm. (F–G) Quantification of the percentages of embryo patterning defects at the 8- to 16-cell stage (F) and embryo arrest (G) in the indicated genotypes. Data are presented as mean ± SD from three biological replicates. Different letters indicate statistically significant differences among groups, as determined by one-way ANOVA followed by Tukey’s multiple-comparisons test (*P* < 0.05).

To further validate the interaction, we performed Co-IP assays using transgenic plants expressing *pMTA:mMTA-GFP* together with *pSWI3B:SWI3B-FLAG*. *MTA-GFP* together with *SWI3B-FLAG* served as a positive control. SWI3B-FLAG co-immunoprecipitated with wild-type MTA-GFP but not with mMTA-GFP. This finding confirmed that amino acid residues K443 and R446 are essential for MTA–SWI3B interaction (Figure 4C).

To further investigate the functional significance of this interaction, we examined whether *pMTA:mMTA* could complement the embryonic defects of *mta-2*. Despite screening multiple transformants, homozygous *pMTA:mMTA mta-2* plants were not discovered. Siliques of *pMTA:mMTA mta-2/+* contained approximately 17% white ovules with arrested embryos in two independent transgenic lines, whereas the *pMTA:MTA* transgene fully rescued embryo lethality in *mta-2* (Figures 4D, 4G). Similarly, approximately 16% of *pMTA:mMTA mta-2/+* embryos displayed patterning defects at the 16-cell stage, while these defects were substantially rescued by *pMTA:MTA* (Figures 4E, 4F). Together, these results demonstrate that the interaction between MTA and SWI3B is required for the developmental function of MTA during embryogenesis.

### MTA and SWI3B are required for proper apical-basal axis formation and auxin maxima during early embryo patterning

Next, we asked whether MTA and SWI3B regulate embryo patterning by controlling key embryonic regulators. The transcription factors WOX2 and WOX8 specify the apical and basal embryonic lineages, respectively (Ueda, et al., 2011; Breuninger, et al., 2008). To examine apical-basal axis formation, we introduced the dual reporter line *pWOX2:NLS-DsRed/ pWOX8:NLS-YFP* into *mta-2/+* and *swi3b-2/+* plants. In the segregating progeny of *mta-2/+*, about one-quarter (24.1%, n=29) of embryos exhibited reduced *WOX8* expression in the hypophysis and the upper one or two suspensor cells (Figure 5A, middle panels). A similar phenotype was observed in about one-quarter of *swi3b-2/+* embryos (23.5%, n = 34), in which *WOX8* expression was likewise reduced in the hypophysis and upper suspensor cells. This indicates altered basal cell identity (Figure 5A, lower panels). In contrast, *WOX2* expression pattern was indistinguishable from that of the wild type in both mutants (Figure 5A).

**Figure 5.**
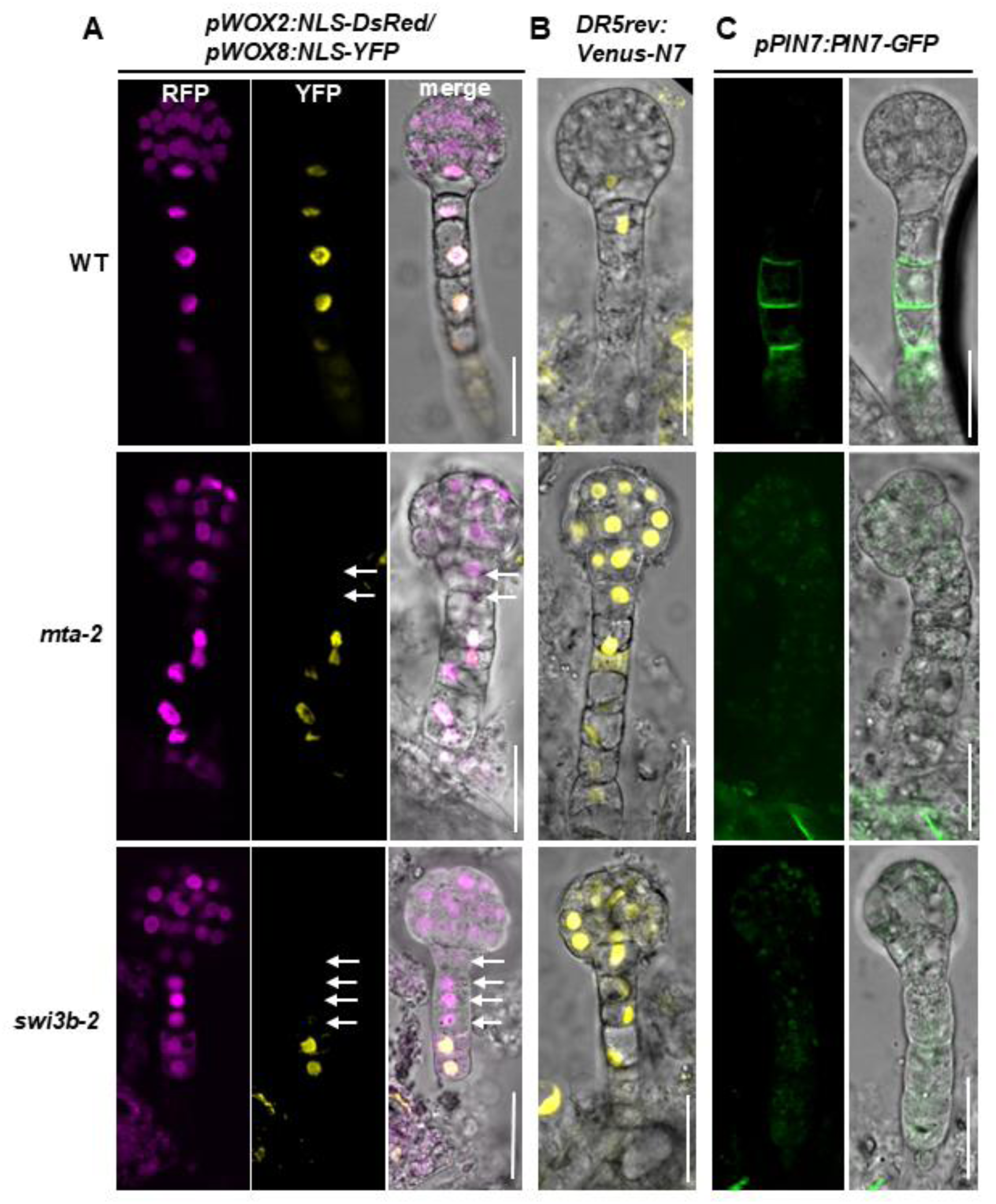
MTA and SWI3B are required for proper apical-basal axis formation and auxin maxima during early embryo patterning. (A) Expression patterns of *pWOX2:NLS-DsRed*/*pWOX8:NLS-YFP* in globular-stage embryos of WT, *mta-2*, and *swi3b-2*. White arrows indicate loss of *WOX8* expression in the hypophysis and suspensor cells. (B) Expression pattern of *DR5rev:Venus-N7* in globular-stage embryos of WT, *mta-2*, and *swi3b-2*. (C) Expression pattern of *pPIN7:PIN7-GFP* in 16- to 32-cell-stage embryos of WT, *mta-2*, and *swi3b-2*. Scale bars, 20 μm.

Because auxin signaling is essential for early embryo patterning (Friml, et al., 2003), we next examined auxin response and transport. At the early globular embryo stage, the auxin response reporter *DR5rev* marked auxin maxima in the hypophysis and the uppermost suspensor cell in wild-type embryos (Figure 5B, upper panel). By contrast, approximately one-quarter of *mta-2/+* (24.4%, n = 45) and *swi3b-2/+* (26.5%, n = 49) embryos displayed expanded and increased *DR5rev* signals throughout the embryo, coinciding with abnormal cell division patterns and morphology in these regions (Figure 5B, middle and lower panels). We next analyzed the polar auxin efflux carrier PIN7, which is required to establish the apical-basal axis and auxin maxima during early embryogenesis (Friml, et al., 2003). In wild-type early globular embryos, PIN7-GFP showed polar localization in suspensor cells (Figure 5C, upper panels). In the phenotypic fraction of *mta-2/+* (26.0%, n = 23) and *swi3b-2/+* (25.9%, n = 27) embryos, PIN7-GFP exhibited cytoplasmic accumulation, indicative of impaired polar localization (Figure 5C, middle and lower panels). Together, these results demonstrate that MTA and SWI3B are required for proper embryo patterning by maintaining *WOX8* expression, PIN7 localization, and auxin distribution during early embryogenesis.

### SWI3B, MTA, and FIP37 share a subset of target transcripts during embryogenesis

To investigate the molecular mechanisms by which SWI3B and the m^6^A writer complex regulate gene expression during embryogenesis and to identify downstream targets, we isolated phenotypic pre-globular embryos from segregating *swi3b-2/+*, *mta-2/+*, and *fip37-4/+* siliques (presumptive homozygous mutant embryos) together with wild-type embryos and performed RNA-seq (Figure 6A). Principal component analysis (PCA) and hierarchical clustering demonstrated high reproducibility among biological replicates and clear separation of the four genotypes (Figures 6B and S6). Among the mutants, *swi3b-2* embryos showed the greatest transcriptional divergence from the wild type, whereas *mta-2* displayed an intermediate profile and *fip37-4* clustered closer to the wild type, consistent with the relative severity of their embryonic phenotypes at this stage. Differential expression analysis identified 1,205, 772, and 327 differentially expressed genes (DEGs) in *swi3b-2*, *mta-2*, and *fip37-4* embryos, respectively, compared with the wild type (|log2FoldChange| ≥ 1, adjusted P < 0.05) (Figure 6D). Notably, several of the most strongly upregulated genes in *swi3b-2* were also significantly upregulated in *mta-2* and *fip37-4* (Figure 6D, circled). Among the upregulated genes, 90 were shared among all three mutants (Figure 6C). In contrast, only 15 downregulated genes were common to all three mutants (Figure S7B). These data suggest that SWI3B and the m^6^A writer complex regulate a common set of downstream genes.

**Figure 6.**
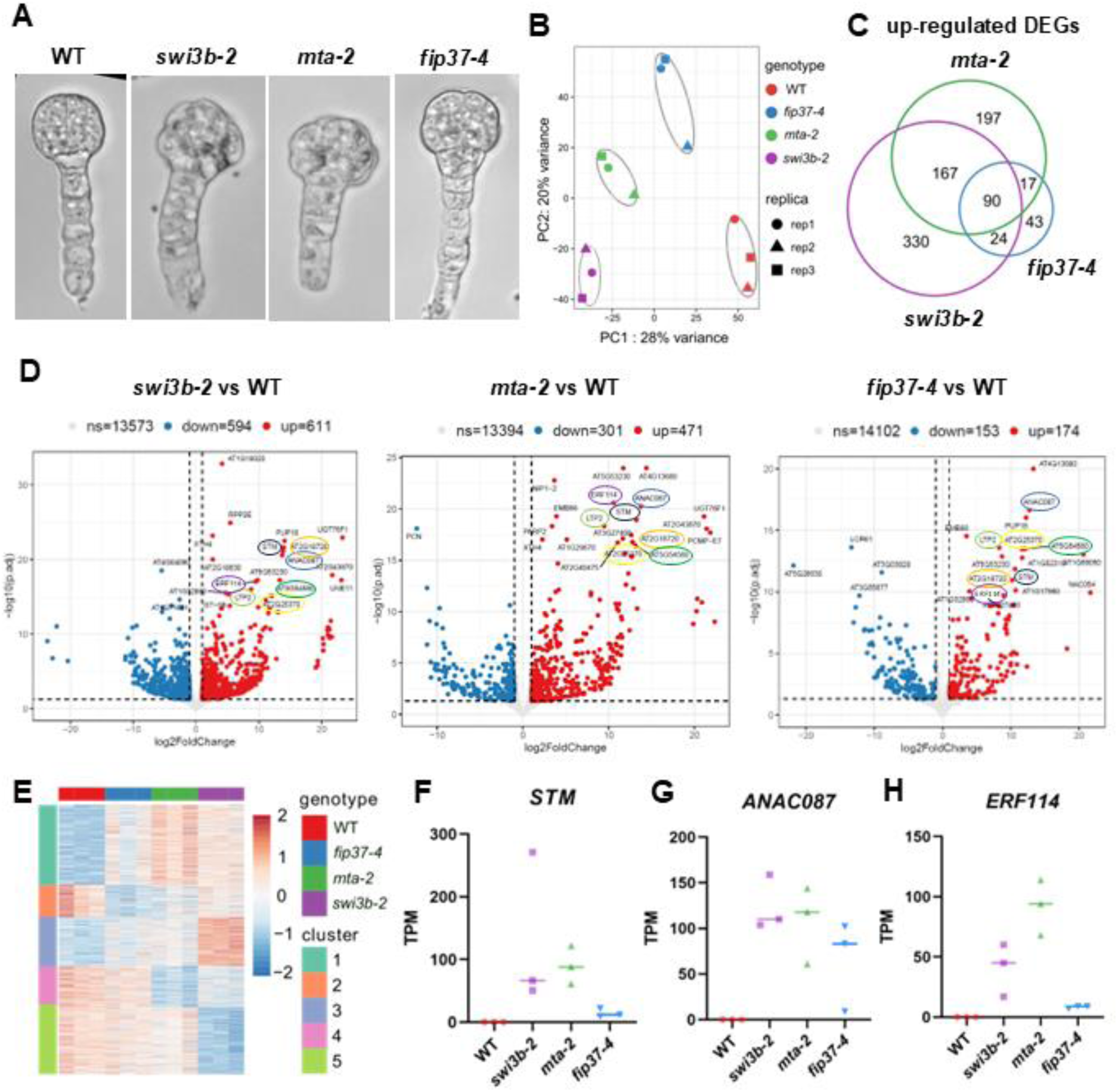
SWI3B, MTA, and FIP37 regulate a common subset of transcripts during embryogenesis. (A) DIC microscopy of isolated 32-cell-stage embryos of WT, *swi3b-2*, *mta-2*, and *fip37-4*. (B) Principal component analysis (PCA) of RNA-seq samples showing clustering of biological replicates and separation among genotypes. (C) Venn diagram showing the numbers and overlap of differentially expressed genes (DEGs) in *mta-2*, *fip37-4*, and *swi3b-2* relative to WT embryos. (D) Volcano plots showing up- and down-regulated DEGs in the three mutants relative to WT. Red and blue dots indicate significantly up- and down-regulated genes, respectively. Selected highly significant DEGs are labeled, with commonly up-regulated DEGs highlighted by colored circles. (E) Heatmap of gene expression profiles grouped by k-means clustering. Each column represents one biological replicate of the indicated genotype, and colors indicate row-wise z-scores of gene expression. (F–H) Transcripts per million (TPM) values of *STM* (F), *ANAC087* (G), and *ERF114* (H) in three RNA-seq biological replicates of WT, *swi3b-2*, *mta-2*, and *fip37-4* embryos.

To identify genes exhibiting similar expression patterns across the three mutants, we performed k-means clustering of all detected genes. This analysis identified five gene expression clusters (Figure 6E). Cluster 1 contained genes that were up-regulated in all three mutants, with the strongest induction in *mta-2*. In contrast, no cluster exhibited consistent down-regulation in all three mutants. Cluster 2 contained genes down-regulated in *mta-2* and *fip37-4*, whereas cluster 5 contained genes down-regulated in *swi3b-2* (Figure 6E, S7A). To determine whether the commonly up-regulated genes are associated with specific biological processes, we performed Gene Ontology (GO) enrichment analysis on Clusters 1 and 5. Both clusters were significantly enriched for genes involved in transcriptional regulation, while Cluster 1 was additionally enriched for senescence-related processes (Figure S7C). We therefore focused our subsequent analyses on transcription factors that were commonly upregulated in all three mutants (Table 1). Among the most strongly induced genes were *SHOOT MERISTEMLESS* (*STM*), *ANAC087*, *ANAC085*, *HDG4*, *ERF114*, *ERF014*, and *WRKY57*. Strikingly, these transcripts were barely detectable in wild-type pre-globular embryos but accumulated to high levels in all three mutant backgrounds.

**Table 1.** Topmost upregulated genes in three mutants.

|  | WT | *swi3b-2* | *mta-2* | *fip37-4* |
| --- | --- | --- | --- | --- |
| PUP18 | 0 | 191.4625 | 32.73148 | 77.3972 |
| AT2G18720 | 0 | 115.9873 | 61.21816 | 10.28303 |
| STM | 0 | 270.8089 | 88.46721 | 12.01513 |
| ANAC087 | 0 | 110.4477 | 117.8597 | 83.21937 |
| AT5G54560 | 0 | 117.0006 | 139.8213 | 48.88569 |
| AT2G25370 | 0 | 47.5789 | 54.62778 | 49.34939 |
| AT1G52315 | 0 | 27.06942 | 58.91938 | 53.02318 |
| AT4G13680 | 0 | 20.73867 | 323.4962 | 137.1832 |
| AT1G21830 | 0 | 40.96887 | 28.66598 | 4.289149 |
| AT1G17960 | 0 | 44.36377 | 150.6326 | 62.76649 |
| HDG4 | 0 | 35.83987 | 23.11784 | 5.700446 |
| ANAC085 | 0 | 15.3921 | 37.56522 | 17.63592 |
| CDS3 | 0 | 10.3762 | 26.48741 | 11.36112 |
| AT2G40475 | 0 | 56.41741 | 65.97958 | 6.802827 |
| AT5G53230 | 0 | 27.958 | 255.5944 | 28.44631 |
| ARAB-1 | 0 | 4.306498 | 39.41725 | 7.94473 |
| ERF114 | 0.117647 | 59.63906 | 94.14084 | 8.601295 |
| AT3G27400 | 0.062281 | 35.87853 | 84.28684 | 9.516124 |
| LTP2 | 0 | 231.9749 | 361.0799 | 151.3219 |
| AGO3 | 0 | 26.07507 | 19.72493 | 11.59348 |
| AT1G27110 | 0 | 4.510842 | 11.01103 | 8.616318 |
| SUMO4 | 0 | 8.129635 | 18.55412 | 37.90309 |
| GRP3 | 0.046433 | 111.3955 | 264.4855 | 325.0952 |
| WRKY57 | 0.501677 | 28.57328 | 23.54276 | 15.48369 |
| ERF014 | 0.795764 | 39.21436 | 81.9949 | 68.77826 |

*STM* has previously been identified as a direct target of FIP37-mediated m^6^A modification (Shen, et al., 2016), suggesting that m^6^A-dependent regulation of *STM* transcript abundance also operates during embryogenesis (Figure 6F). In addition, ANAC087, a positive regulator of programmed cell death (Huysmans, et al., 2018), and ERF114, a regeneration-associated transcription factor that integrates mechanical cues with auxin signalling (Canher, et al., 2022), were among the most strongly induced genes in all three mutants. Given their known biological functions, we tested whether their ectopic expression may contribute to the developmental arrest, abnormal embryo morphology, and altered auxin responses observed in the mutants. Consistent with this possibility, *ANAC087* and *ERF114* transcripts were virtually undetectable in wild-type embryos but accumulated extensively in the mutant embryos (Figures 6G and 6H).

### Elevated expression of ANAC087 and ERF114 during embryogenesis leads to embryo defects

We therefore studied whether ectopic accumulation of *STM*, *ANAC087* and *ERF114* transcripts contributes to the embryo patterning defects observed in *swi3b-2*, *mta-2* and *fip37-4* mutants. To test this possibility, we generated constructs expressing *STM*, *ANAC087* and *ERF114* under the early embryo-specific promoter *pCZ2* (Zhao, et al., 2019). Despite extensive screening of T1 seedlings, we were unable to recover any *pCZ2:STM* transformants. We assume that elevated *STM* expression during early embryogenesis severely impairs embryo development. For *pCZ2:ANAC087* and *pCZ2:ERF114*, two independent transformant lines were obtained for each construct. Approximately 20–25% of embryos exhibited abnormal cell division patterns in the suspensor of these lines (Figures 7A and 7D). Further analysis revealed defects in embryo proper morphology and aberrant hypophysis cell divisions. These phenotypes resembled those observed in *swi3b-2*, *mta-2*, and *fip37-4* embryos (Figure 7B). Consistently, at later developmental stages, siliques containing green mature embryos also contained a similar proportion of white ovules, corresponding to arrested embryos (Figures 7C and 7E). As a negative control, *pCZ2:GFP* transgenic plants did not exhibit any embryo patterning defects or arrest (Figure 7A-E). These results indicate that ectopic expression of *STM*, *ANAC087* and *ERF114* during embryogenesis is sufficient to induce embryo developmental or even lethal defects.

**Figure 7.**
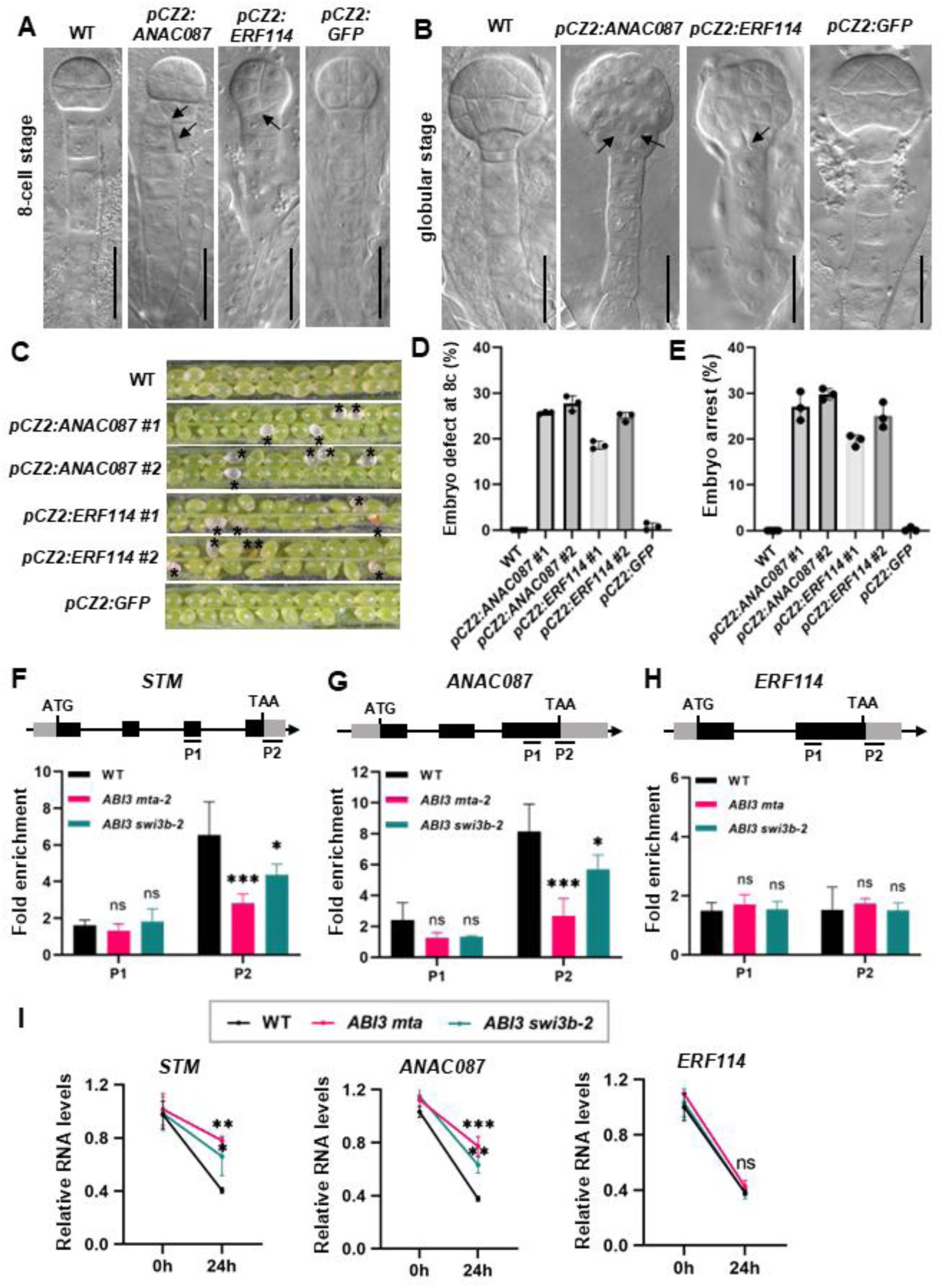
Elevated levels of *ANAC087* and *ERF114* mRNAs during embryogenesis lead to embryo defects. (A-B) DIC microscopy of 8-cell stage (A) and globular stage (B) embryos in WT, *pCZ2:ANAC087*, *pCZ2:ERF114* and *pCZ2:GFP* plants. Arrows point to abnormal cell division planes. Bars, 20 μm. (C) Dissected siliques from indicated genotypes. Asterisks indicate white aborted ovules. (D-E) Quantification of the percentages of embryo patterning defects at 8-cell stage (D) and embryo arrest (E) in indicated genotypes. Data are presented as mean ± SD from three biological replicates. (F-H) m^6^A modification levels in *STM* and *ANAC087* mRNAs are compromised in *pABI3:MTA mta* and *pABI3:SWI3B swi3b-2* in five-day-old seedlings. m^6^A-IP-qPCR showed m^6^A modification levels in *STM* (F), *ANAC087* (G) and *ERF114* (H) mRNA, respectively. Upper panels show transcript structure including cDNA fragments amplified in qPCR (P1 and P2). Error bars, mean ± SD from three biological replicates. Each biological replicate corresponds to three technical replicates. Asterisks indicate statistically significant differences between mutants and the WT by two-way ANOVA with Tukey’s multiple comparisons test (ns, no significance; *, *p* < 0.05; ***, *p* < 0.001). (I) The stability of *STM* and A*NAC087* transcripts is increased in *pABI3:MTA mta* and *pABI3:SWI3B swi3b-2.* qRT-PCR shows the transcript levels of *STM*, *ANAC087* and *ERF114* in 5-day-old seedlings of WT and mutants treated by actinomycin D versus mock for 0 and 24 hours. Relative expression of each gene at each time point was calculated by normalizing the gene expression in actinomycin D treated samples against the mock treated ones. Error bars, mean ± SD from three biological replicates. Each biological replicate corresponds to three technical replicates. Asterisks indicate statistically significant differences between mutants and the WT by Two-way ANOVA with Tukey’s multiple comparisons test (ns, no significance; *, *p* < 0.05; ***, *p* < 0.001).

We next investigated whether SWI3B and MTA regulate transcript abundance of *ANAC087* and *ERF114* through effects on mRNA stability and m^6^A modification. Specifically, we asked whether the stability and m^6^A modification levels of *ANAC087* and *ERF114* transcripts are altered in the absence of SWI3B or MTA. Because homozygous *swi3b* mutants are embryo lethal, we generated a homozygous *swi3b* line in which SWI3B expression is restricted to embryogenesis, thereby allowing analysis of SWI3B function after embryo development. The *ABI3* promoter has previously been shown to rescue the embryo-lethal phenotype of *mta* mutants (*pABI3:MTA mta*, hereafter referred to as *ABI3 mta*) and is active primarily during embryogenesis (Bodi, et al., 2012). We therefore expressed the *SWI3B* coding sequence under control of the *ABI3* promoter and introduced this construct into *swi3b-2/+* plants. Homozygous *pABI3:SWI3B swi3b-2* plants (hereafter referred to as *ABI3 swi3b-2*) were recovered, demonstrating that the construct complemented the embryo-lethal phenotype. Both the early embryo patterning defects and the subsequent embryonic arrest were fully rescued in *ABI3 swi3b-2* plants (Figure S8A-C). RT-qPCR analysis confirmed that *SWI3B* transcript levels were significantly reduced in seedlings and flowers of *ABI3 swi3b-2* plants compared with the WT (Figure S8D). To determine whether SWI3B influences MTA-dependent m^6^A deposition, we performed m^6^A-IP-qPCR using mRNA isolated from 5-day-old WT, *ABI3 mta*, and *ABI3 swi3b-2* seedlings. Enrichment of m^6^A at the 3′ UTR of *STM* was significantly reduced in both, *ABI3 mta* and *ABI3 swi3b-2* (Figure 7F), consistent with previous reports identifying *STM* as an m^6^A-modified transcript (Shen, et al., 2016). Likewise, m^6^A enrichment at the 3′ UTR of *ANAC087* was significantly reduced in both mutants (Figure 7G). In contrast, a significant difference in m^6^A enrichment was not detected in *ERF114* transcripts (Figure 7H). These results indicate that both MTA and SWI3B are required for efficient m^6^A deposition on the 3′ UTRs of *STM* and *ANAC087*.

As m^6^A has been implicated in promoting mRNA decay in mammalian systems (Zlotorynski, 2025; Wei, 2024). We therefore finally examined whether reduced m^6^A deposition affects the stability of *STM* and *ANAC087* transcripts. Five-day-old WT, *ABI3 mta*, and *ABI3 swi3b-2* seedlings were treated with the transcriptional inhibitor actinomycin D, and transcript abundance was measured 24 h after treatment relative to mock-treated controls. Both *STM* and *ANAC087* transcripts exhibited significantly greater stability in *ABI3 mta* and *ABI3 swi3b-2* compared with the WT, whereas the stability of *ERF114* transcripts was not significantly altered (Figure 7I). Altogether, these results indicate that SWI3B promotes MTA-dependent m^6^A deposition on *STM* and *ANAC087* mRNAs, thereby facilitating the degradation of these transcripts.

## Discussion

Our findings identify the SWI/SNF chromatin-remodeling complex subunit SWI3B as a functional interactor of the m^6^A writer MTA and reveal a role for this interaction in the regulation of early Arabidopsis embryogenesis. Several lines of evidence support a functional connection between SWI3B and the m^6^A writer complex. *swi3b-2* and *mta-2* embryos exhibited similar early patterning defects, including altered auxin distribution and reduced *WOX8* expression in the suspensor, and transcriptome analysis revealed a substantial group of genes commonly deregulated in *swi3b-2*, *mta-2*, and *fip37-4* embryos. Importantly, disruption of the SWI3B– MTA interaction impaired the developmental function of MTA without abolishing its interaction with FIP37, providing evidence that the association with SWI3B contributes specifically to MTA function. Furthermore, several transcription factors that are normally expressed at very low levels during early embryogenesis accumulated strongly in all three mutants, and ectopic expression of selected candidates was sufficient to perturb embryo patterning. Together, these findings establish a previously unrecognized connection between SWI/SNF-dependent chromatin regulation and the m^6^A machinery and suggest that SWI3B contributes to selective RNA regulation by the MTA-containing writer complex during embryogenesis.

Despite their overlapping functions, SWI3B and MTA make distinct contributions to embryo development. The phenotype of *swi3b-2* was more severe than that of *mta-2*: *swi3b-2* embryos arrested predominantly at the pre-globular stage, whereas *mta-2* embryos underwent several additional rounds of cell division after the initial patterning defects before arresting. Consistent with this difference, substantially more differentially expressed genes were detected in pre-globular *swi3b-2* embryos than in *mta-2* embryos. This broader transcriptional effect is consistent with the central role of SWI3B as a core component of SWI/SNF chromatin-remodeling complexes and suggests that only part of its developmental function is mediated through its association with the m^6^A writer complex. SWI3B also functions at other stages of reproductive development. In particular, we recently found that SWI3B is required for the initiation of megagametogenesis by restricting *AGO5* expression in the ovule nucellus through regulation of the local chromatin environment (Gong et al., bioRxiv, 2026). Thus, SWI3B appears to regulate multiple transitions during plant reproduction through chromatin-dependent control of developmental gene expression, whereas its interaction with MTA may provide an additional layer of regulation for a subset of transcripts during embryogenesis.

MTA and FIP37 likewise showed overlapping but non-identical embryo developmental phenotypes and transcriptome profiles. Both proteins are core components of the plant m^6^A writer complex, and loss of either protein causes embryonic lethality. However, the early embryonic phenotype of *fip37-4* was less pronounced than that of *mta-2*, making mutant embryos more difficult to distinguish morphologically from their wild-type siblings and consequently limiting reporter-based analyses. Consistent with this milder early phenotype, transcriptome analysis identified substantially fewer differentially expressed genes in *fip37-4* than in *mta-2* embryos. These differences may reflect distinct molecular contributions of individual writer-complex components or differences in the timing and severity of their developmental consequences. Notably, studies in mammalian and Arabidopsis have identified functions of METTL3/MTA that can be uncoupled from its canonical activity within the m^6^A writer complex (Liu, et al., 2021; Bhat, et al., 2020; Choe, et al., 2018; Lin, et al., 2016), raising the possibility that individual plant writer components may likewise have additional or partially independent functions. Whether MTA or FIP37 has such functions in plants remains to be determined. Importantly, despite these differences, the common transcriptional signature of *swi3b-2*, *mta-2*, and *fip37-4* identifies a set of genes whose repression during early embryogenesis depends on the shared activities of SWI3B and the m^6^A machinery.

The early patterning defects observed in *swi3b-2* and *mta-2* were accompanied by alterations in several key components of the apical–basal patterning network, including changes in *DR5rev* auxin response, PIN7 distribution, and reduced *WOX8* expression in the suspensor. However, whether these changes contribute directly to the initiation of the patterning defects or arise as subsequent consequences of abnormal embryo development remains unclear. In particular, changes in *DR5rev* and PIN7–GFP signals could only be detected from approximately the 16-cell stage onwards with our microscopy setup, whereas abnormal cell-division patterns were already evident at earlier stages. Nevertheless, perturbation of PIN7- and WOX8-dependent pathways can produce auxin-patterning defects resembling those observed in *swi3b-2* and *mta-2*. PIN7 is the earliest PIN auxin efflux carrier detected during embryogenesis, with endogenous protein detectable by immunolocalization from the one-cell stage, although reporter-based detection does not fully recapitulate this early endogenous expression pattern (Moller and Weijers, 2009; Vieten, et al., 2005; Friml, et al., 2003). Loss of *PIN7* results in increased auxin response in the suspensor (Robert, et al., 2013), while *wox8 wox9* double mutant similarly exhibits enhanced auxin response together with severe early embryo-patterning defects and embryonic lethality (Breuninger, et al., 2008). Thus, impaired PIN7 polarity and reduced *WOX8* expression could contribute to the altered auxin distribution and subsequent developmental defects observed in *swi3b-2* and *mta-2*. However, given the temporal relationship between the reporter phenotypes and the onset of abnormal cell divisions, our data do not distinguish whether disruption of these pathways represents a primary consequence of impaired SWI3B–MTA function or a secondary amplification of earlier patterning defects.

Our transcriptome and functional analyses further identify developmental regulators that may contribute to the mutant phenotypes. Several transcription factor genes were virtually undetectable in wild-type pre-globular embryos but accumulated to high levels in *swi3b-2*, *mta-2*, and *fip37-4*. Embryo-specific expression of *STM* did not yield transformants, consistent with the possibility that inappropriate *STM* expression during early embryogenesis severely compromises development. Although *STM* is expressed in shoot apical meristem stem cells from the transition stage onwards, its absence during earlier stages suggests that precise temporal control of *STM* expression is essential for normal embryo development (Balkunde, et al., 2017). Moreover, embryo-specific expression of *ANAC087* or *ERF114* was sufficient to phenocopy the abnormal cell-division phenotypes observed in the mutants, demonstrating that deregulation of individual members of this transcriptional program can have substantial developmental consequences. ANAC087 is a positive regulator of developmental programmed cell death (PCD) and is sufficient to activate PCD-associated genes and induce ectopic cell death when misexpressed (Chen, et al., 2023; Huysmans, et al., 2018). PCD is itself an integral part of normal embryo development: the suspensor is a transient structure that supports the embryo during early development and is subsequently eliminated through PCD (Peng and Sun, 2018; Luo, et al., 2016; Blanvillain, et al., 2011). Although a role for ANAC087 in suspensor PCD has not been established, these observations emphasize the importance of restricting the activity of potent PCD regulators both spatially and temporally during embryo development. Our finding that *ANAC087* transcripts accumulate prematurely in *swi3b-2*, *mta-2*, and *fip37-4* embryos, together with the embryo defects caused by ectopic *ANAC087* expression, raises the possibility that m^6^A-dependent RNA turnover contributes to this temporal restriction by preventing inappropriate accumulation of *ANAC087* transcripts during early embryogenesis. Our data further suggest that the deregulated genes arise through at least two mechanistically distinct downstream branches. *STM* and *ANAC087* represent candidate direct targets of m^6^A-dependent regulation: their transcript abundance increases when the m^6^A machinery is disrupted. By contrast, the available evidence does not support *ERF114* as a direct m⁶A-regulated transcript, suggesting that its accumulation may result from an m⁶A-independent function of SWI3B–MTA or from indirect downstream effects.

How SWI3B contributes to the selectivity of this RNA-regulatory pathway remains unresolved. One possibility is that chromatin-associated SWI3B helps recruit or position the MTA-containing writer complex at particular genomic loci through its physical interaction with MTA, thereby promoting m⁶A deposition on selected nascent transcripts. Alternatively, SWI3B-dependent chromatin remodeling may establish a local chromatin environment permissive for efficient m⁶A deposition, while the SWI3B–MTA interaction stabilizes or positions the writer complex at these loci. These possibilities are not mutually exclusive and are consistent with evidence from mammalian systems that chromatin states and chromatin-associated factors can influence co-transcriptional m⁶A deposition (Hu, et al., 2024; Wei, 2024). Our finding that disruption of the SWI3B–MTA interaction compromises MTA function provides a functional basis for such a model, although direct recruitment of MTA to SWI3B-bound chromatin remains to be demonstrated.

## Limitations of the study

Several limitations should be considered when interpreting our findings. Owing to the limited amount of material obtainable from early embryos, m^6^A enrichment and transcript stability were assessed in five-day-old seedlings rather than directly in embryos. Whether the same m^6^A-dependent regulatory events occur during embryogenesis therefore remains to be established. Moreover, transcript stability was assessed at a single time point after 24 h of actinomycin D treatment rather than by determining complete RNA decay kinetics, and prolonged transcriptional inhibition may introduce indirect or stress-related effects. Although increased transcript abundance of *STM*, *ANAC087*, and *ERF114* was detected in the mutants, corresponding changes in protein abundance could not be examined because of the limited amount of embryonic material. In addition, embryo-specific overexpression of *ANAC087* and *ERF114* phenocopied aspects of mutant defects but does not demonstrate that endogenous de-repression of these genes is necessary for the *swi3b-2* or *mta-2* phenotypes. Finally, because a transcriptome-wide analysis of m^6^A deposition was not feasible with the available embryonic material, we cannot determine whether SWI3B broadly influences the embryonic m^6^A landscape or regulates a restricted subset of MTA targets.

## Resource availability

### Lead contact

Requests for further information and resources should be directed to and will be fulfilled by the lead contact, Wen Gong.

## Materials availability

This study did not generate new unique reagents.

## Data availability

All sequencing data have been deposited at GEO as GSE342816 and are publicly available as of the date of publication.

## Acknowledgments

We thank Armin Hildebrand for plant care, Maria Hutterer for cloning and genotyping work, Tomasz Sarnowski for kindly providing us with the mutant seeds of *swi3b-2/+*, and Zofia Szweykowska-Kulinska for the seeds of *pABI3:MTA mta*. The German Research Foundation (DFG) is acknowledged for financial support via SFB960 (to TD).

## Author contributions

WG and TD initiated the project. WG designed and performed the experiments. US performed bioinformatic analysis. LF performed the protein structure prediction. GL supported for protein-RNA binding assays. TD provided research funding and input. WG and TD wrote the manuscript with input from all authors.

## Declaration of interests

None

## Declaration of generative AI and AI-assisted technologies in the writing process

During manuscript preparation, we used ChatGPT to improve language clarity and readability. All content was carefully reviewed and edited by the authors, who take full responsibility for the final manuscript.

## Materials and Methods

### Materials and growth conditions

*Arabidopsis thaliana* (L.) Heynh. (Arabidopsis) ecotype Columbia (Col-0; wild type, WT), mutant, and reporter lines were grown on soil under long-day conditions (16 h light at 8500 lux, 21°C, and 65% humidity). The mutants used in this study are *swi3b-2/+* (GABI_302G08), *mta-2/+* (SALK_123823) and *fip37-4* (SALK_018636). Primers used for genotyping PCRs are listed in Table S1.

### Plasmid construction

Primers used for cloning are listed in Table S1. For *pMTA:MTA-*GFP, 2035-bp upstream of the start codon of *MTA* was taken as the promoter, 3363-bp genomic region from the start codon until the sequence before the stop codon was amplified. For *pFIP37:FIP37-*GFP, 1538-bp upstream of the start codon of *FIP37* was taken as the promoter, 2546-bp genomic region from the start codon until the sequence before the stop codon was amplified. For *pSWI3B:SWI3B-GFP* and *pSWI3B:SWI3B-mScarlet*, 2029-bp upstream of the start codon of *SWI3B* was taken as the promoter, 2124-bp genomic region after the promoter before the stop codon was amplified. For Co-IP assay, the whole genomic regions including the promoters of *MTA*, *FIP37* and *SWI3B* without stop codon were amplified. A C-terminal 1x FLAG tag coding sequence (Einhauer and Jungbauer, 2001) was introduced by PCR amplification. For the GFP, mScarlet and FLAG fusion constructs, a GSAGAG-linker was added before the tags (Chen, et al., 2013). For the negative control *pGIF2:GIF2-FLAG*, 3595-bp genomic region of *GIF2* (AT1G01160) before the stop codon was amplified together with the GSAGAG-linker and cloned together with a FLAG tag. For *pMTA:MTA*, the whole genomic region of *MTA* (5401-bp) was amplified. For *pMTA:mMTA* and *pMTA:mMTA-GFP*, the point mutations were generated by PCR using the primers containing the mismatches (a1327g, c1336g, g1337a and t1338g), using *pMTA:MTA* and *pMTA:MTA-GFP* as the template respectively. For *pCZ2:ANAC087*, *pCZ2:ERF114* and *pCZ2:STM*, 2585-bp upstream of *MYB62* (AT1G68320) was taken as the promoter with additional 21 bp of the first exon of *MYB62*. The coding sequences (CDS) of *ANAC087*, *ERF114* and *STM* were amplified from Arabidopsis cDNA from pistils. For *pABI3:SWI3B*, 4962-bp upstream of the *ABI3* start codon was used as the promoter. The *SWI3B* CDS was amplified from Arabidopsis cDNA using pistils. Amplified fragments were assembled using NEBuilder HiFi DNA Assembly Master Mix (New England Biolabs) to the vector *pGreenII0125* containing a *35S* terminator. The resistance is kanamycin in *Escherichia coli* and norflurazon *in planta*. For BiFC assays, the CDSs of *MTA* and *SWI3B* were amplified and ligated into pRT-SPYNE, a vector containing 35S:nYFP. The CDSs of *FIP37*, *SWI3B* and *GIF2* were amplified and ligated into pRT-SPYCE, a vector carrying 35S:cYFP (Walter, et al., 2004).

### Microscopy

Phenotypic analysis of embryo development was made by using differential interference contrast (DIC) microscopy (Imager.A2, Zeiss). Siliques at different developmental stages were dissected, followed by transferring of ovules connected by the septum to a drop of clearing solution (chloral-hydrate: H₂O:glycerol = 8:2:1 solution). After the tissues were cleared, the samples were observed under the DIC microscope using either a 40x or a 63x objective.

The expression pattern of reporter lines was observed using a Leica DMi8 spinning-disk confocal microscope with a Visitron system equipped with an HC PL APO 63×/1.40-0.60 oil objective. Siliques at different developmental stages were dissected, followed by transferring of ovules connected by the septum to a drop of 10% glycerol. After placing the cover slip, samples were pressed with a round bottom pen on the cover slip, to get the embryos squeezed out of the ovules. YFP and Venus were excited at 505 nm, RFP was excited at 561 nm, and GFP was excited at 488 nm.

### Bimolecular fluorescence complementation (BiFC)

For BiFC assays, Arabidopsis leaf mesophyll protoplasts were isolated and transfected for transient gene expression as previously described (Wu, et al., 2009; Yoo, et al., 2007). After overnight incubation in darkness following transfection with respective plasmids, fluorescence signals from protoplasts were observed under previously described Leica DMi8 spinning-disc confocal laser microscope with an HC PL APO 40×/1.3 NA water objective. YFP was excited at 505 nm, and chlorophyll was excited at 561 nm.

### Co-Immunoprecipitation (Co-IP)

For Co-IP assays, 5-day-old transgenic seedlings carrying indicated constructs were harvested for IP. *pMTA:MTA-GFP / pSWI3B:SWI3B-FLAG* was used as the F1 generation from a cross between *pMTA:MTA-GFP* and *pSWI3B:SWI3B-FLAG*. The same was done for *pMTA:MTA-GFP / pGIF2:GIF2-FLAG*, *pMTA:MTA-GFP / pFIP37:FIP37-FLAG* and *pMTA:mMTA-GFP / pSWI3B:SWI3B-FLAG*, which are the F1 generations from corresponding crosses. About 1 g of 5-day-old seedlings per sample was ground in liquid nitrogen with a mortar and pestle. The manufacturer’s protocol for the GFP-Trap agarose kit (Chromotek) was followed for IP. 10% of the samples were saved as the input samples before adding agarose beads and immediately boiled for 5 min at 95 °C with the equal volume of 2x SDS-sample buffer. After protein binding and washing, 80 μL 2x SDS-sample buffer was added and boiled for 5 min at 95 °C to elute proteins. Supernatants of IP samples and input samples were analyzed by Western blots. For Western blots, inputs and immunoprecipitated samples were loaded on 10% SDS-PAGE gels. Then, blots were probed with anti-GFP (Abcam, ab290) or anti-FLAG (Proteintech, 20543-1-AP), each with a 1:5000 dilution. Next blots were probed with the second antibody HRP-goat anti-rabbit (Proteintech, RGAR001) with a 1:5000 dilution. Blots were detected by chemiluminescent substrate (Thermo Fisher Scientific, A38555) and the ChemiDoc MP Imaging system (Bio-Rad) for chemiluminescence.

### m^6^A-IP (MeRIP) qPCR

m^6^A IP was performed using 5-day-old seedlings of WT, *ABI3 mta* and *ABI3 swi3b-*2 grown on MS plates. Total RNA was extracted using the RNeasy Plus Mini kit (Qiagen) according to the manufacturer’s instructions. The EpiQuik Cut&Run m^6^A RNA Enrichment (MeRIP) kit (EpigenTek, P-9018) was used for m^6^A-IP, RNA fragmentation and purification, following the manufacturer’s instructions. 20 μg total RNA was used per IP. 200 ng RNA was used per input. Purified RNA samples were reverse transcribed with random hexamers (Invitrogen) using ReverseAid RT (Thermo Fisher Scientific). Relative enrichment of each gene was determined by quantitative real-time PCR (qPCR). For each target, qPCR was performed with two primer pairs targeting the exon and the 3’ UTR. The primers are listed in Table S1. qPCR was performed using the Eppendorf ep realplex Mastercycler real-time PCR system and the SYBR Green Master Mix (Roche). Three biological replicates and three technical replicates for each sample were performed in the m^6^A-IP qPCR experiments. For each sample and each primer pair, m^6^A enrichment level was calculated as RQ (relative quantity) of IP/RQ of Input.

### mRNA stability analysis

5-day-old seedlings of WT, *ABI3 mta* and *ABI3 swi3b-2* grown on MS plates were transferred to liquid MS medium containing 10 μM actinomycin D (Sigma) or DMSO as previously described (Shen, et al., 2016). Samples were harvested at the beginning of the treatment (0h) and 24 hours after the treatments (24h). Total RNA was extracted using the RNeasy Plus Mini kit (QIAGEN) according to the manufacturer’s instructions, and reverse transcription performed using ReverseAid RT (Thermo Fisher Scientific) with 1 μg total RNA. qPCR was performed with the primers specifically targeting the cDNA of *STM*, *ANAC087* and *ERF114*. The primers for qRT-PCR are listed in Table S1. The relative mRNA level was calculated as RQ of actinomycin D-treated/RQ of mock-treated.

### Embryo isolation

Embryo isolation was carried out following a previously described method (Zhao, et al., 2019) with slight modifications. Self-pollinated siliques were harvested for collecting ovules for embryo isolation. Siliques were dissected under a stereoscopic microscope, and ovules were transferred to 100 μl of enzyme solution in the bottom of a 3.5-cm Petri dish. Ovules were treated with cell-wall-degrading enzyme solution for 30 min at room temperature. Next, the enzyme solution was removed, and 100 μl of washing solution was added to wash ovules three times. Globular stage embryos were dissected directly from ovules with two fine needles under an inverted microscope (Eclipse TS100, Nikon). Isolated embryos were transferred to another droplet of washing solution by a handmade capillary pipette, then transferred to 10 μl of RNAlater (Thermo Fisher Scientific), and kept at −80°C for later usage. For segregating mutant embryos, only defective embryos were isolated. About 20 isolated embryos were each pooled as one replicate for RNA-seq.

### Total RNA extraction and mRNAseq

Total RNA was extracted from isolated embryos and stabilized in RLT Plus buffer according to the protocol of the RNeasy Plus Micro Kit (Qiagen). In brief, cells were stored in 350 μl of buffer RLT Plus containing 1% ß-mercaptoethanol and shipped on dry ice. After thawing, samples were homogenized by vortexing for 1 min. Genomic DNA contamination was removed by using gDNA Eliminator spin columns (Sigma-Aldrich). Next, one volume of 70% ethanol was added, and each sample was applied to RNeasy MinElute spin columns (Qiagen) followed by several washing steps. Finally, total RNA was eluted in 12 μl of nuclease-free water. Purity and integrity of RNA were assessed using the Agilent 2100 Bioanalyzer with the RNA 6000 Pico LabChip reagent set (Agilent).

Library preparation and mRNAseq were carried out as described in the SMART-Seq mRNA LP User Manual (Takara Bio), the Illumina NextSeq 2000 Sequencing System Guide (Illumina Inc.), and the KAPA Library Quantification Kit – Illumina/ABI Prism Protocol (Roche Sequencing Solutions Inc.). In brief, around 2 ng of total RNA was used to generate first-strand cDNA. Double-stranded cDNA was amplified by LD-PCR (12 cycles) and purified via magnetic bead cleanup. After validating the quality and quantity, about 150 pg of cDNA was enzymatically fragmented and stem-loop adapters were ligated. Then, ligated fragments were PCR-amplified (16 cycles) and indexed, generating Illumina-compatible libraries with unique dual indexes. After a magnetic bead purification, libraries were quantified using the KAPA Library Quantification Kit (Roche). Equimolar amounts of each library were sequenced on an Illumina NextSeq 2000 instrument controlled by the NextSeq 2000 Control Software v.1.5.0.42699 using a 100 cycles P3 Flow Cell with the dual index, single-read run parameters. Image analysis and base calling were made by the Real Time Analysis Software v.3.10.30. The resulting .cbcl files were converted into .fastq files with the bcl2fastq v.2.20 software (www.illumina.com/company/legal.html).

### RNA-seq analysis

Unstranded single-end RNA-seq data were annotated using an in-house nextflow RNA-seq pipeline (Wernig-Zorc, et al., 2024; Di Tommaso, et al., 2017) (available at GitHub: https://github.com/uschwartz/RNAseq_NAC; v2.1). Initially, quality control of the raw sequence reads was conducted using FastQC (v0.11.8) (https://www.bioinformatics.babraham.ac.uk/projects/fastqc/4). Adapter sequences were trimmed from the 3’ ends using Trim Galore (v0.6.7) (https://github.com/FelixKrueger/TrimGalore?tab=readme-ov-file). Reads were mapped to the Arabidopsis reference genome (TAIR10) and the corresponding gene annotation (Ensembl version 47) using Spliced Transcripts Alignment to a Reference (STAR v2.7.8a) (Dobin, et al., 2013). The following options were used to optimize the alignment process: --outFilterType BySJout, --outFilterMultimapNmax 20, --alignSJoverhangMin 8, --alignSJDBoverhangMin 1, - -outFilterMismatchNmax 999, --alignIntronMin 10, --alignIntronMax 1,000,000, -- outFilterMismatchNoverReadLmax 0.04, --runThreadN 12, --outSAMtype BAM SortedByCoordinate, --outSAMmultNmax 1, and --outMultimapperOrder Random.

Postmapping quality control was performed using the rnaseq analysis mode of Qualimap (v2.2.1) (García-Alcalde, et al., 2012). The level of PCR duplication was assessed using Picard MarkDuplicates (v2.21.8) (https://github.com/broadinstitute/picard) and dupRadar (v1.15.0) (Sayols, Scherzinger and Klein, 2016). Gene expression quantification for all annotated genes was carried out using featureCounts (v1.6.3) (Sayols, et al., 2016).

### Differential gene expression analysis

Pairwise differential gene expression analysis of *swi3b-2*, *mta-2*, and *fip37-4* samples against WT samples were performed using the Bioconductor DESeq2 package (Love, et al., 2014). Prior to statistical testing genes with a TPM of less than one in more than three samples were removed and further filtered for protein-coding and lncRNA biotypes. The ashr shrinkage method was applied to properly scale log2 (fold-changes) (Stephens, 2017). Differentially expressed genes (DEGs) were identified using an FDR of 0.05 and a |log2(fold-change)|>1 as cutoffs.

### k-means clustering and functional gene enrichment analysis

For clustering all DEGs in any pairwise comparison against WT controls were considered. The median rlog value across each genotype was extracted for each gene. Next, the median rlog values were scaled to z-scores across all genotypes and split into five distinct clusters using k-means clustering. The genes were split according to cluster association and multiple gene lists gene ontology (GO) analysis against a background of all expressed genes was performed in Metascape (Zhou, et al., 2019). Here, a p-value of 0.0001 and a minimum overlap of 5 genes were used as cutoffs.

## Supplemental information

**Figure S1.**
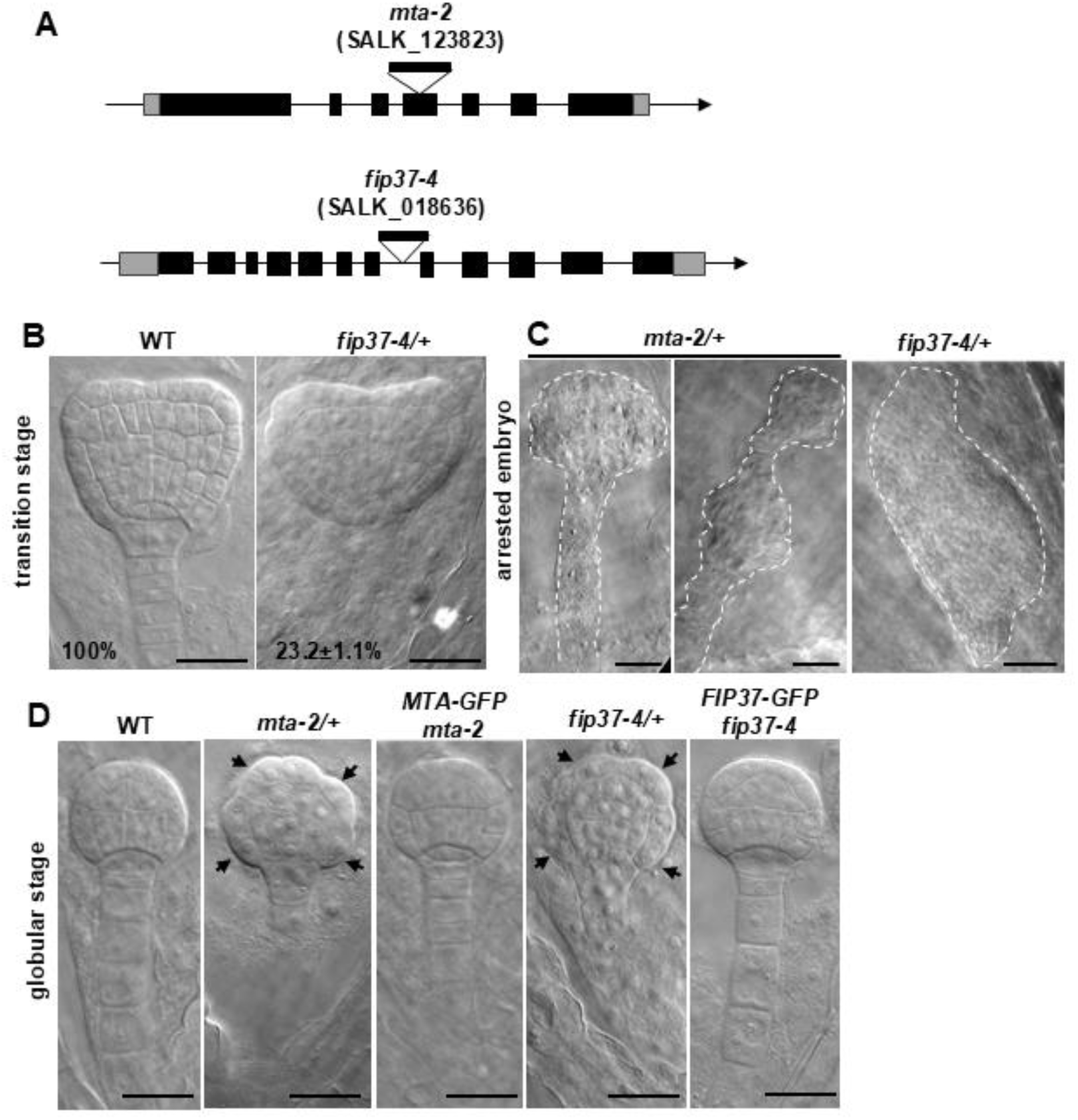
Characterization of *mta-2* and *fip37-4* embryo phenotypes and complementation lines. (A) Schematic representation of the genomic regions of *MTA* and *FIP37*. T-DNA insertion sites and the corresponding SALK accession numbers are indicated. Black boxes indicate exons, and gray boxes indicate 5′ and 3′ UTRs, respectively. (B) DIC microscopy of transition-stage embryos of WT and *fip37-4/+*. Percentages of the indicated phenotypes are shown. Scale bars, 20 μm. (C) DIC microscopy of arrested embryos from white ovules of *mta-2/+* and *fip37-4/+*. Dashed lines indicate embryo outlines. Scale bars, 20 μm. (D) DIC microscopy of globular-stage embryos of WT, *mta-2*, *pMTA:MTA-GFP mta-2*, *fip37-4/+*, and *pFIP37:FIP37-GFP fip37-4*. Scale bars, 20 μm.

**Figure S2.**
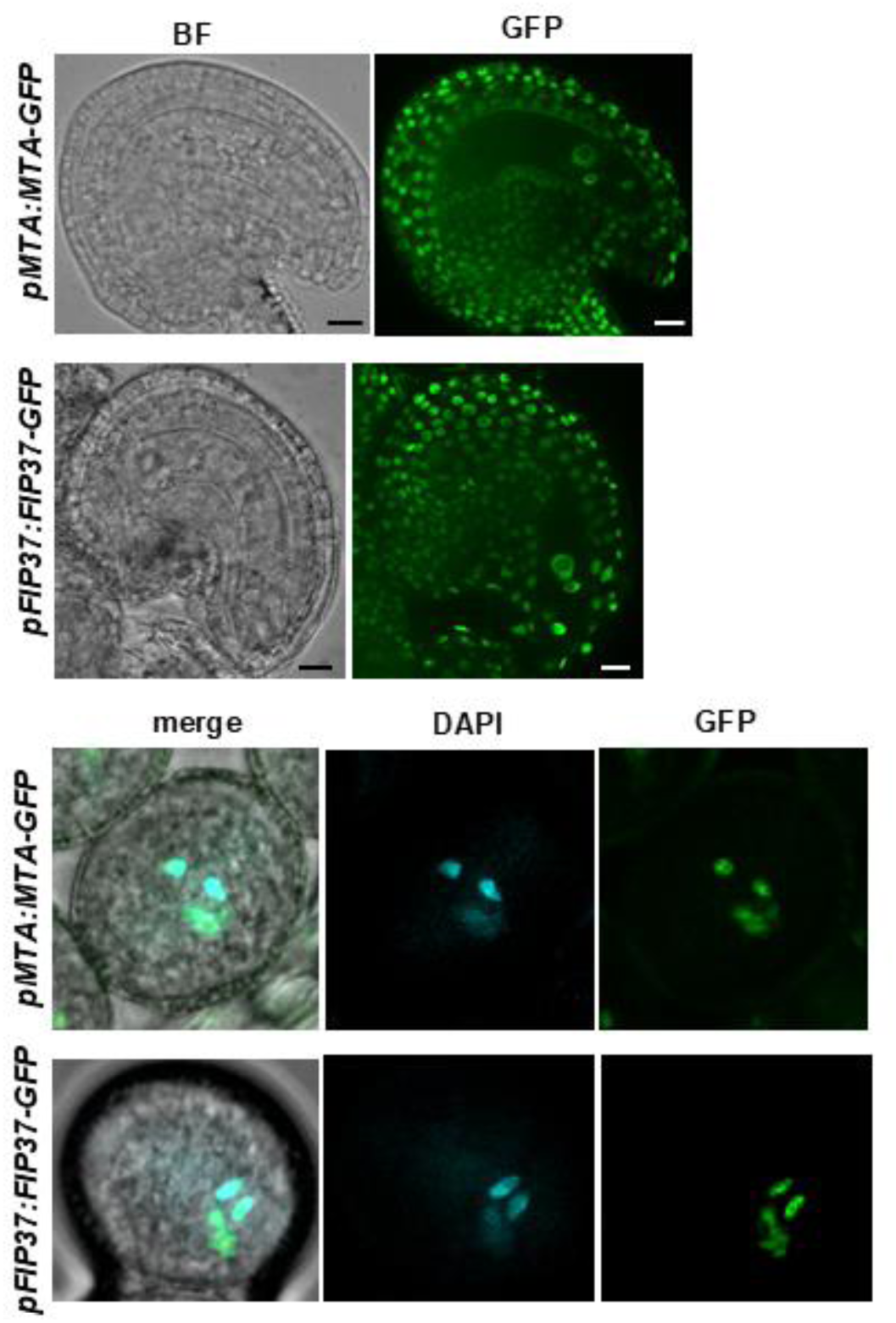
MTA and FIP37 are expressed in male and female gametes. Expression patterns of *pMTA:MTA-GFP* and *pFIP37:FIP37-GFP* in unfertilized ovules (upper panels) and pollen grains (lower panels) as indicated. Pollen nuclei were stained with DAPI. Scale bars, 20 μm.

**Figure S3.**
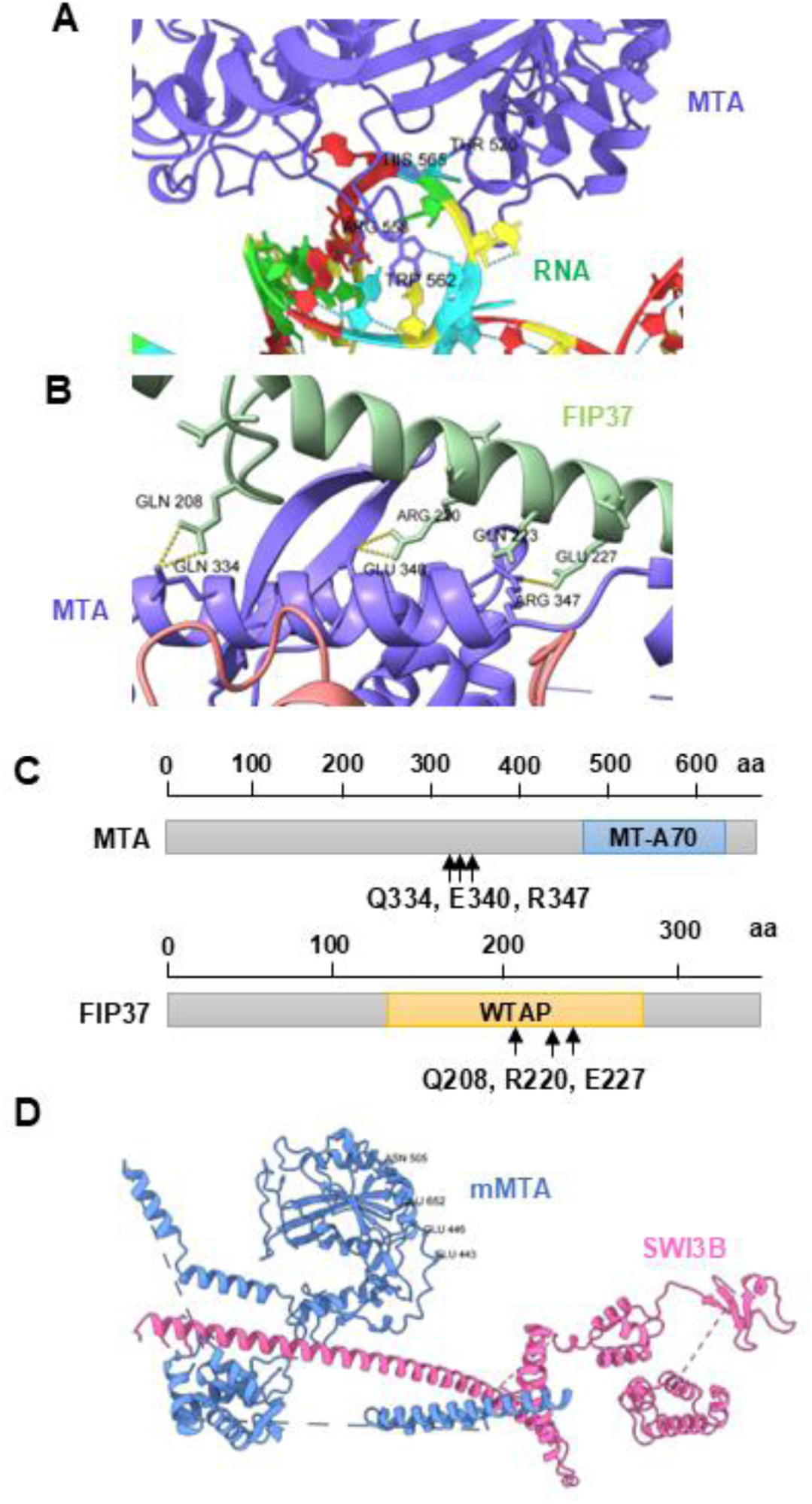
AlphaFold3 modeling predicts MTA residues involved in interactions with FIP37, SWI3B and RNA. (A) AlphaFold3 model of predicted interaction sites between MTA (blue) and RNA. The RNA target is the same as shown in Figure 2D. The ribonucleotides of the RNA are shown in different colors. (B) AlphaFold3 model of the predicted interaction sites between MTA (blue) and FIP37 (green). Amino acid residues predicted to mediate the MTA-FIP37 interaction are indicated. (C) Schematic representation of MTA and FIP37 proteins. Amino acid positions are indicated above schematics, and colored boxes denote functional domains. Arrows indicate amino acid residues predicted to mediate protein interactions. (D) AlphaFold3 model predicting loss of the interaction between mutated MTA (mMTA; blue) and SWI3B (red). Substituted amino acid residues of MTA are indicated.

**Figure S4.**
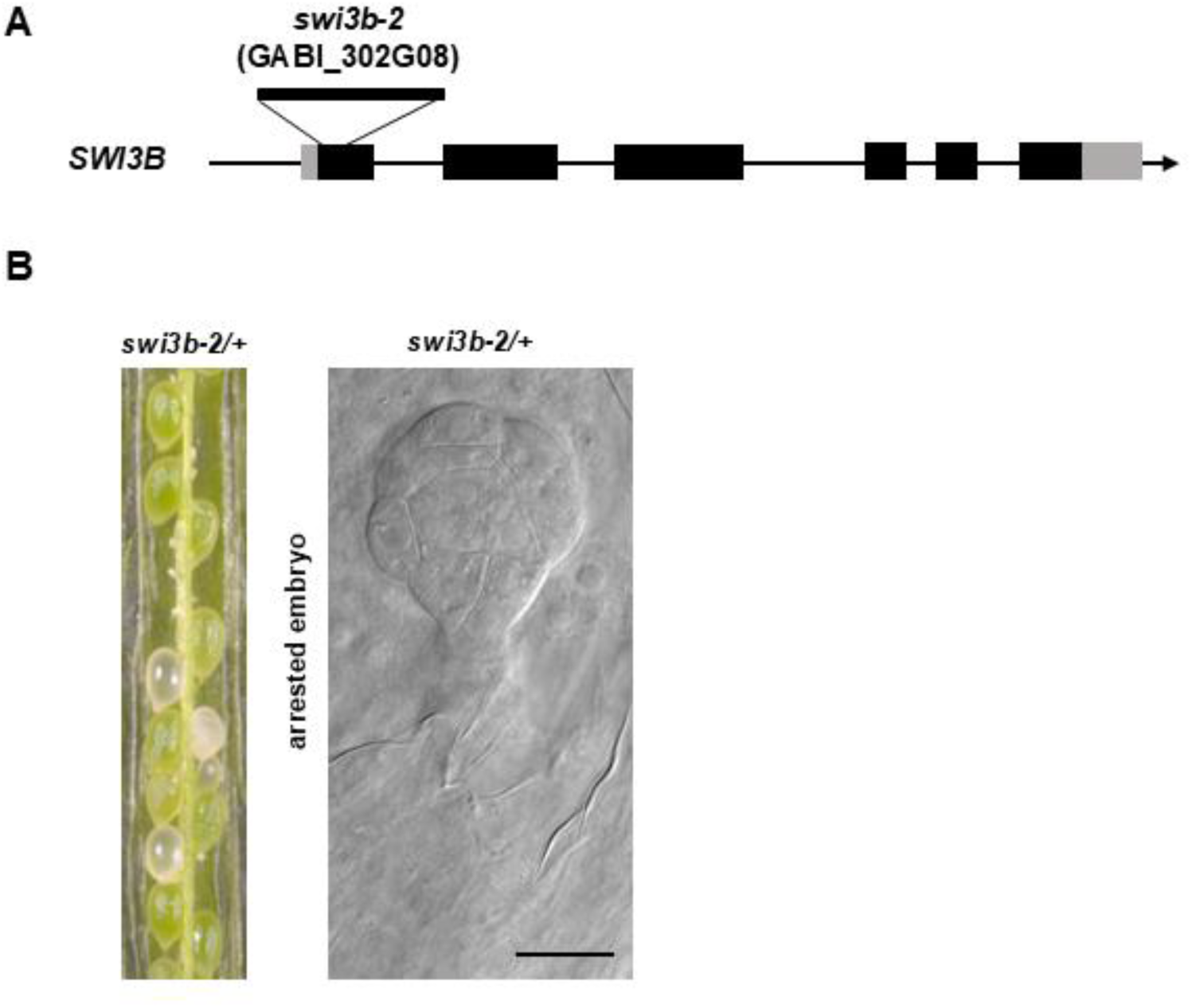
Characterization of the embryo-lethal phenotype of *swi3b-2*. (A) Schematic representation of the genomic region of *SWI3B*. The T-DNA insertion site in *swi3b-2* is indicated. Black boxes indicate exons, and gray boxes indicate 5′ and 3′ UTRs. (B) Dissected silique of *swi3b-2/+* (left) and DIC microscopy of an arrested *swi3b-2* embryo at the early globular stage (right). Scale bar, 20 μm.

**Figure S5.**
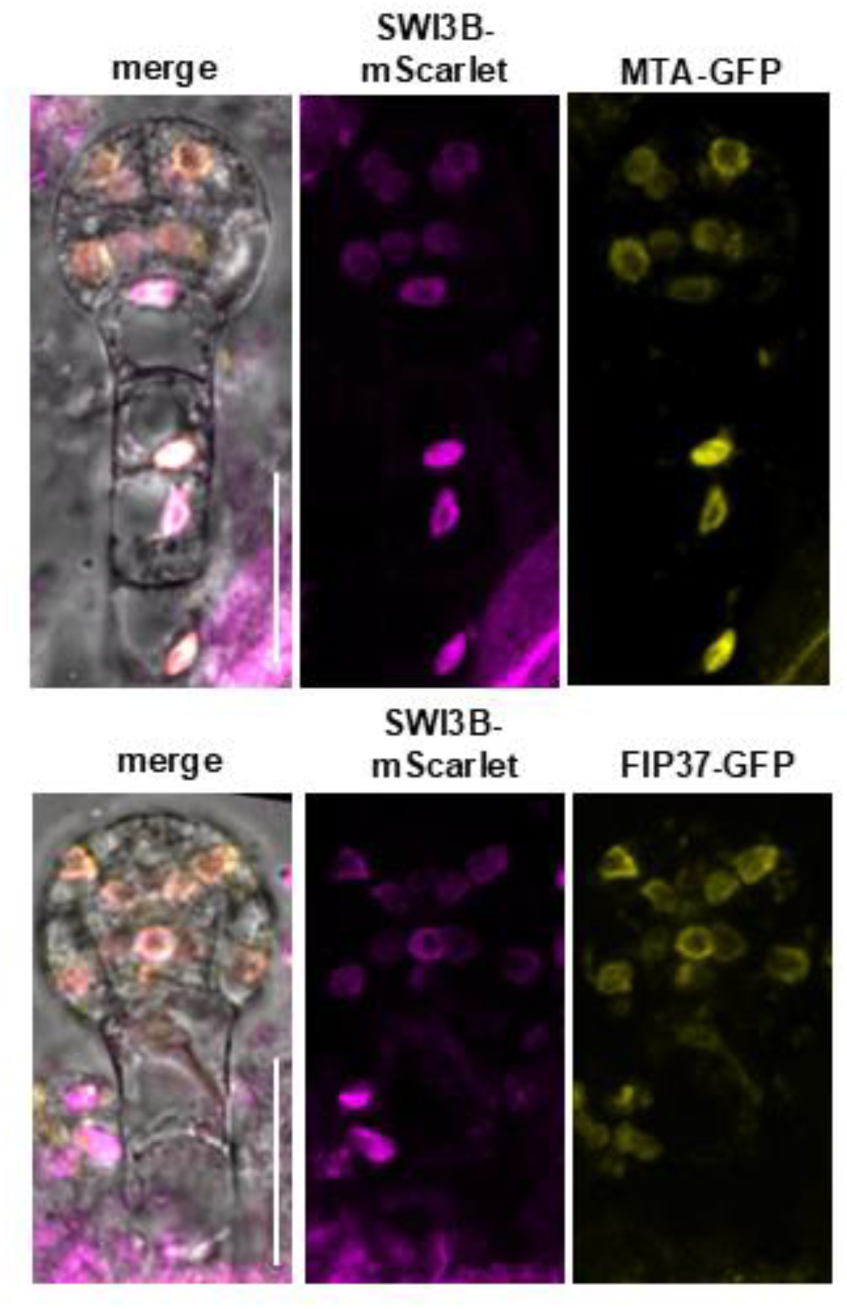
SWI3B colocalizes with MTA and FIP37 in nuclei of pre-globular embryos. Colocalization of MTA-GFP and SWI3B-mScarlet (upper panels) and FIP37-GFP and SWI3B-mScarlet (lower panels) in nuclei of pre-globular-stage embryos. Scale bar, 20 μm.

**Figure S6.**
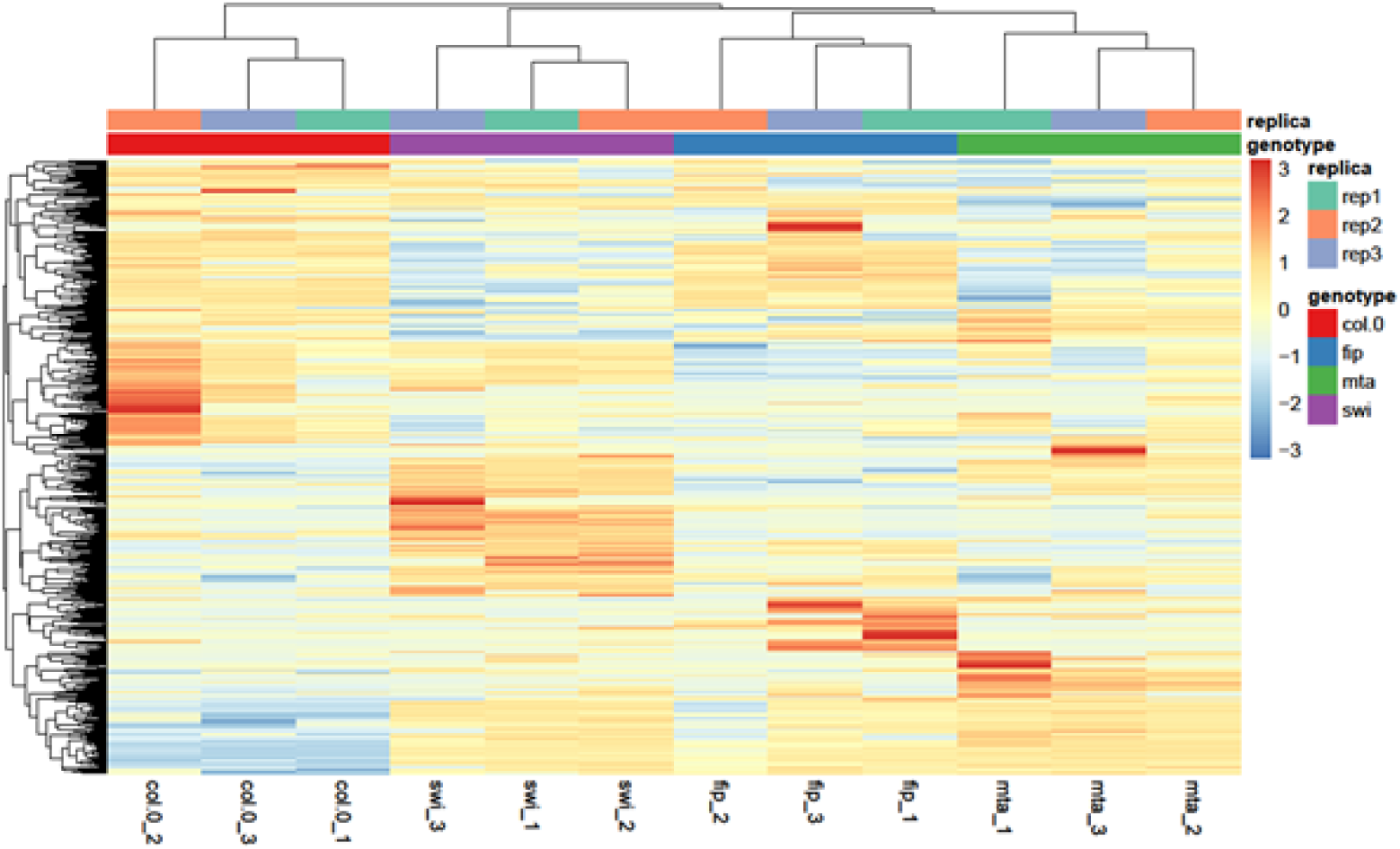
Hierarchical clustering of transcriptomes from WT, *mta-2*, *fip37-4*, and *swi3b-2* embryos. Heatmap showing hierarchical clustering of RNA-seq of biological replicates from WT, *mta-2*, *fip37-4*, and *swi3b-2* embryos. Genotypes and biological replicates are indicated by annotated colors. The heatmap color scale represents row-wise z-scores of relative gene expression.

**Figure S7.**
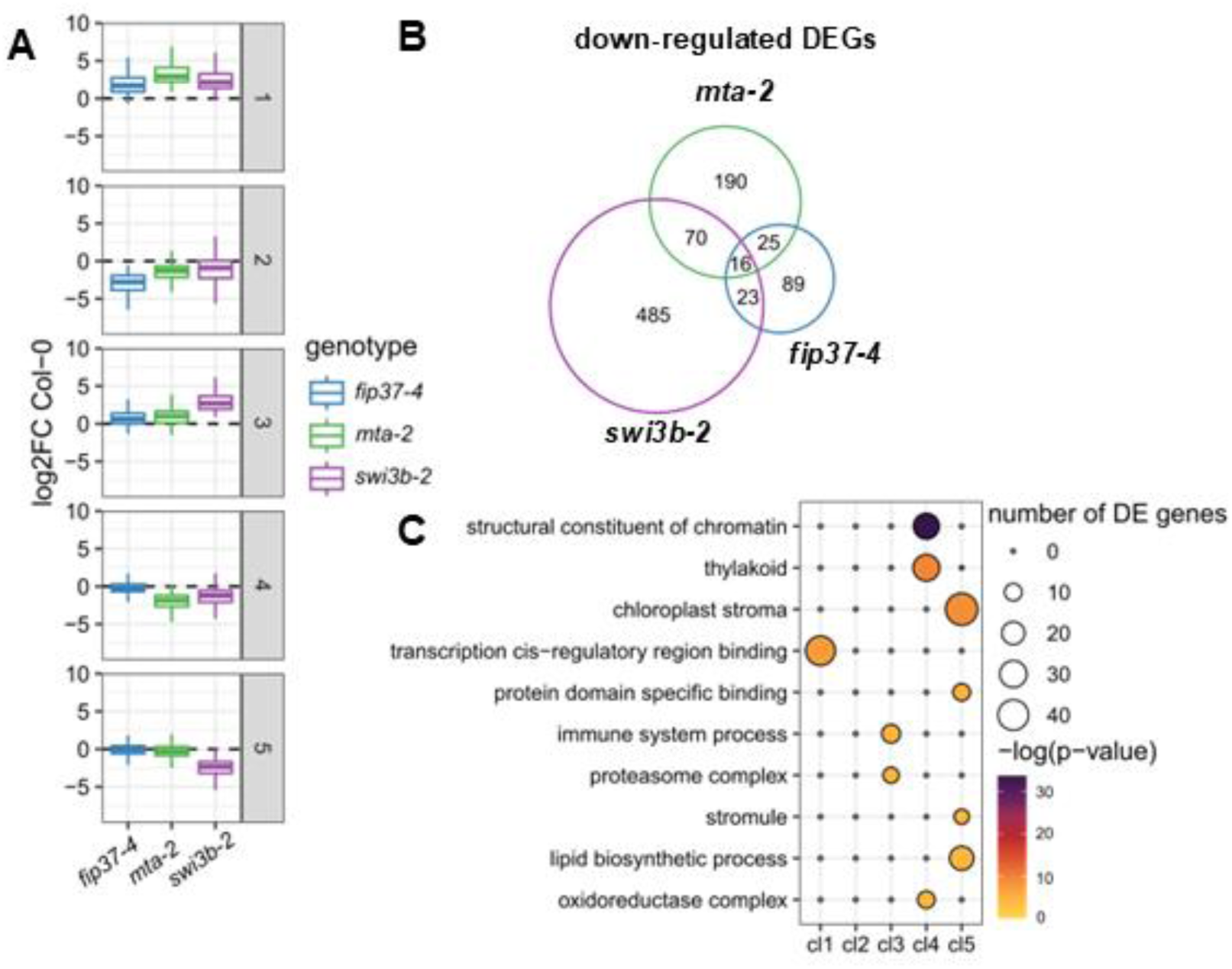
Expression pattern and functional enrichment of DEG clusters. (A) Boxplots showing the average log2 fold changes (log2FC) of genes assigned to the five fuzzy c-means clusters shown in Figure 6E in *fip37-4*, *mta-2*, and *swi3b-2* relative to WT embryos. Center lines indicate medians. Dashed line indicates no change relative to WT embryos (log2FC = 0). (B) Venn diagram showing the numbers and overlap of significantly downregulated DEGs in *fip37-4*, *mta-2*, and *swi3b-2*. (C) Gene Ontology (GO) enrichment analysis of biological process terms for genes assigned to k-means clusters 1-5.

**Figure S8.**
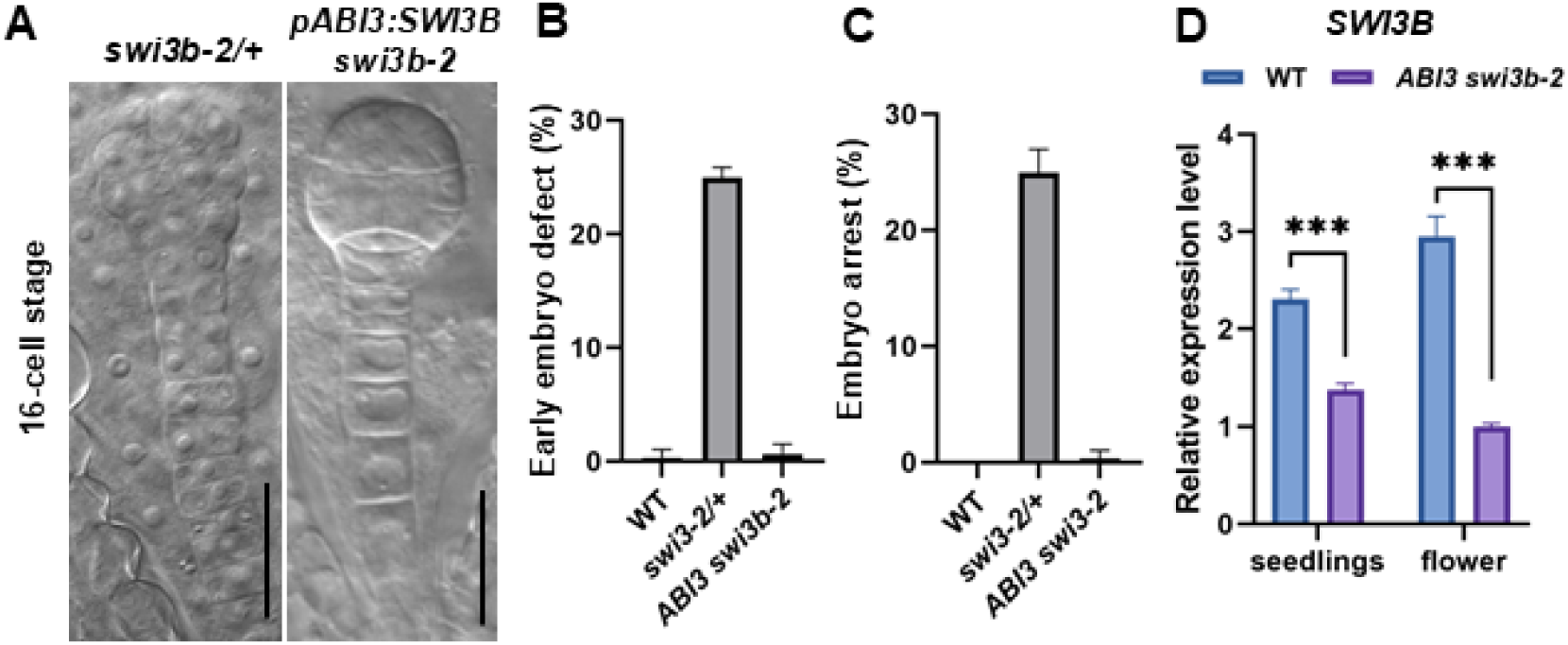
Embryo-specific expression of *SWI3B* complements the embryo defects of *swi3b-2*. (A) DIC microscopy of 16-cell-stage embryos of *swi3b-2/+* and *pABI3:SWI3B swi3b-2*. Scale bars, 20 μm. (B–C) Quantification of the percentages of embryo patterning defects at the 8- to 16-cell stage (B) and embryo arrest (C) in the indicated genotypes. Data are presented as mean ± SD from three biological replicates. (D) Relative *SWI3B* transcript levels in seedlings and flowers of WT and *pABI3:SWI3B swi3b-2*, determined by qRT-PCR. Data are presented as mean ± SD from three biological replicates. Statistical significance was determined by an unpaired two-tailed Student’s *t* test (\*\*\**P* < 0.001).

## Notes

### Competing Interest Statement

The authors have declared no competing interest.

